# Aluminium induces suberin biosynthesis in barley roots via ABA

**DOI:** 10.1101/2024.10.27.620472

**Authors:** Hongjun Meng, Qihui Zhang, Tino Kreszies, Ivan F. Acosta, Lukas Schreiber

## Abstract

Aluminum (Al) toxicity is a major factor limiting plant growth in acidic soils. The beneficial element silicon (Si) can mitigate some effects of Al. However, the impact of Al on suberized apoplastic barriers in roots are largely unknown while the effects of Si on suberin remains controversial.

This study employed physiological, histochemical, and analytical methods, along with Laser Capture Microdissection (LCM) RNA-sequencing, to explore the effects of Al and Si on suberin development in barley (*Hordeum vulgare* L.), a species sensitive to Al stress.

Exposure of barley seedlings to Al resulted in increased suberin deposition, which could be restored with the addition of Si, particularly in the root endodermis. Gene expression analyses using LCM RNA-seq across different root tissues demonstrated that Al-induced suberin biosynthesis is mainly regulated by the abscisic acid (ABA) pathway. In addition, the application of fluridone, an inhibitor of ABA synthesis and a suberin mutant, further supported the pivotal role of ABA in the Al response and the role of suberin in influencing Al uptake.

Our findings underscore the complex interplay between Al stress and suberin biosynthesis in barley, providing insights into potential strategies for enhancing crop resilience to Al toxicity.

## Introduction

Aluminium (Al) is the third most abundant element in the Earth’s crust at 8.2%, while silicon (Si) is the second most abundant, at 27.7% (Exley, 1998). They are primarily present in the soil in the form of insoluble aluminosilicates and their oxides. For most plants, Si is usually considered a beneficial plant nutrient rather than essential (Coskun *et al*., 2019). On the other hand, for most plants, Al is toxic, and it is a crucial factor in acidic soils (pH < 5), in which the Al complexed in aluminosilicate clays is released as the most phytotoxic trivalent cation, Al^3+^ (Kochian, 1995; Kochian *et al*., 2004), often seriously limiting plant growth.

It has been shown that there are many potential Al binding sites in plant cells such as cell walls, cell membranes, and cytoskeleton. Therefore, Al interferes with a wide range of physical and cellular processes (Kochian *et al*., 2005). In most plant species, two main types of Al resistance mechanisms have been described: Al exclusion mechanisms, which prevent Al from entering the root apex, involve Al-induced exudation of organic acid anions and phosphate, including malate, citrate, and oxalate efflux from roots (Kinraide *et al*., 2005; Delhaize *et al*., 2007; Kochian *et al*., 2015; Li *et al*., 2017); and Al tolerance mechanisms, which detoxify and sequester Al in plants, such as fixing Al to the cell wall (Yang *et al*., 2008; Yang *et al*., 2011; Lou *et al*., 2020), sequestering Al into vacuoles for final detoxification (Shen *et al*., 2002; Huang *et al*., 2012), and increasing antioxidant enzyme activity (Wang & Yang, 2005; Zhu *et al*., 2018). In addition, many studies have shown that under certain circumstances soluble Si could ameliorate the toxic effects of Al in many plant species (Hodson & Evans, 1995; Liang *et al*., 2001; Hodson & Evans, 2020).

However, whether the primary damage caused by Al toxicity originates primarily from the apoplast or symplast remains a matter of debate (Zheng & Yang, 2005; Ma, 2007). Al has a strong affinity to electron donors; hence it may target multiple sites simultaneously including negatively charged pectin (Eticha *et al*., 2005; Yang *et al*., 2008) and uncharged hemicellulose (Yang *et al*., 2011; Zhu *et al*., 2012). Here we specifically focus on the effect of Al on suberin, a specialized cell wall component. Suberin in root cell walls plays wide roles in biotic and abiotic stress responses, such as controlling the transport of water and nutrients and limiting the invasion of pathogens (Enstone *et al*., 2003; Franke & Schreiber, 2007; Krishnamurthy *et al*., 2009; Barberon *et al*., 2016; Liska *et al*., 2016; Barberon, 2017; Kreszies *et al*., 2018). In addition, with exposure to some heavy metals, Si and Ca have been found to mediate the development of the suberin lamella, and further affect the absorption and accumulation of heavy metals by plants (Fleck *et al*., 2011; Wu *et al*., 2019; Liu *et al*., 2020). However, whether and how Al affects the development of suberin is still unclear.

Tolerance to Al toxicity or acidic soils differs greatly among cereal crops such as barley (*Hordeum vulgare L*.), which is usually considered one of the cereals most sensitive to Al (Ishikawa *et al*., 2000; Wang *et al*., 2006). Here, we demonstrate that Al stress promotes the development of suberin in barley roots. Subsequent systematic gene expression analyses identified Al-responsive genes in four distinct root tissues: the epidermis, cortex, endodermis, and stele, and revealed a correlation between abscisic acid (ABA) and suberin synthesis. Additionally, our data suggests that suberin affects Al uptake by barley roots.

## Materials and Methods

### Plant material and growth conditions

Two cultivars of barley (*Hordeum vulgare*), Scarlett and Golden Promise Fast (GPF), were used in this study. Seeds were stratified at 4 °C for 2 days, then germinated for 3 days at 25 °C in the dark, and covered with wet filter paper. The 3-day-old (do) seedlings were then transferred into an aerated hydroponic system containing a modified Magnavaca nutrient solution (Famoso *et al*., 2010), and placed in a climatic chamber at 22 °C under a 16 h/8 h light/dark cycle. When plants were 6-do (3 days of germination and 3 days of growth) they were transferred to treatment solutions for another 4 days. For comparisons with CRISPR mutants, *Hordeum vulgare* cv. Golden Promise Fast was used as the normal reference.

### Barley CRISPR mutants

We used the E-CRISP tool (http://www.e-crisp.org/E-CRISP) to choose two sgRNAs targeting the first exon of *CYP86B1* (*HORVU.MOREX.r2.1HG0034810*). The CRISPR-Cas9-sgRNA system was delivered and expressed with Golden Gate vectors kindly provided by Tom Lawrenson and Wendy Harwood (Lawrenson *et al*., 2015) Transformation was carried out in Golden Promise Fast as described (Amanda *et al*., 2022). Mutations in *CYP86B1* were analyzed with PCR and Sanger sequencing. CRISPR sgRNA sequences, cloning primers and gene-specific primers are marked in supplementary table S1.

### Stress application

Six-do plants were transferred to Al treatment solution, which consisted of Magnavaca nutrient solution with 50 μM or 100 μM AlCl_3_, added after pH adjustment to 7.8 with KOH to prevent Al precipitation, and the final pH was adjusted to 4.5 with HCl. Plants grown in Magnavaca nutrient solution at pH 4.5 were used as control. For Si treatment, the modified Magnavaca nutrient solution contained 50 μM or 100 μM AlCl_3_ supplemented with 1 mM Si (NaSiO_3_) was adjusted to pH 4.5 before and after NaSiO_3_ addition. For fluridone treatments, 6-do barley plants (cv. Scarlett) were transferred to Magnavaca nutrient solution containing 10 µM fluridone added after adjusting the medium pH to 4.5. Because fluridone is not directly soluble in water, it was first dissolved in 3 ml DMSO, then 3 ml TWEEN 20 were added, and this solution was diluted to 50 ml with water to obtain a 100 mM fluridone stock solution.

### Chemical analysis of barley root suberin

The 10-do barley seminal roots were divided into three zones, A, B, and C (Kreszies *et al*. 2019). The youngest part of the root (0–25% of total root length), including the root apex, was designated zone A. The middle part of the root (25–50% of total root length) was assigned to zone B. The oldest part of the root (50–100% of total root length), was designated zone C.

For each replicate, at least 10 segments of seminal roots from each of the three zones were pooled together. The root segments were enzymatically digested for 3 weeks with 0.5% (w/v) cellulase and 0.5% (w/v) pectinase at room temperature under continuous shaking (Zeier & Schreiber, 1997). The enzyme solution was replaced four times within the 3 weeks and roots were vacuum infiltrated with the solution. Subsequently, isolated suberized cell walls were washed in borate buffer and then transferred to 1:1 (v/v) chloroform: methanol for soluble lipid extraction at room temperature under continuous shaking for 2 weeks. The chloroform: methanol solution was replaced four times. Finally, samples were dried on polytetrafluoroethylene in a desiccator containing activated silica gel. The dried samples were subjected to transesterification with BF_3_–methanol to release suberin monomers (Kolattukudy & Agrawal, 1974). Gas chromatographic analysis and mass spectrometric identification were performed as described earlier (Zeier & Schreiber, 1997; Zeier & Schreiber, 1998). Suberin amounts were referred to the endodermal surface area. The endodermal area was calculated for each root zone: A = 2π · r · L (r, endodermis radius; L, length of the individual root zone). Three biological replicates were used for each experiment.

### Histochemical staining and microscopy

For suberin staining, seminal roots were cut into pieces of about 1 cm length, which contained the region of interest, and immersed in 4% paraformaldehyde in 1× PBS overnight at 4°C. After fixation, samples were washed three times in 1× PBS. Fixed samples were directly used for clearing with ClearSee solution (10% xylitol, 15% sodium deoxycholate and 25% urea, Kurihara *et al*., 2015) for 5 days at room temperature. Cleared samples were rinsed once in ddH_2_O and immersed in 0.01% Fluorol Yellow 088 (FY) solution for 30 min at room temperature. For cross-sectioning, stained root fragments were embedded in 7% agarose and sectioned by hand with a fresh razorblade. Cross-sections were observed by epifluorescence microscopy with an ultraviolet (UV) filter set (excitation filter BP 365, dichroic mirror FT 395, barrier filter LP 397; Zeiss).

For Al staining, roots were stained in 0.01% (w/v) morin for 30 min, then excised and embedded in 5% (w/v) agar. Root tips were transversely sectioned from the apex, and the green fluorescence signal was observed with laser scanning confocal microscopy (LSM510, Zeiss and Leica SP8 Lightning).

### RNA isolation and RT–qPCR analysis

Total RNA was extracted using the NucleoSpin RNA Plant Mini Kit (MACHEREY-NAGEL, Germany). First-strand cDNA synthesis was performed using the RevertAid First Strand cDNA Synthesis Kit (Thermo Scientific). RT–qPCR was performed in a QuantStudio™ 3 Real Time PCR System (Applied Biosystems) in conjunction with the my-Budget 5x EvaGreen^®^ QPCR-Mix II kit (Bio-Budget Technologies GmbH, Germany). At least three biological replicates of each sample and three technical replicates of each biological replicate were performed to ensure the accuracy of the results. The reference genes *GADPH* or *Actin* were used as internal controls. Primers used for RT–qPCR are listed in supplementary table S1.

### Laser capture dissection microscopy and RNA-seq

An approximate length of 4 mm of root segment originating from the terminal region of Zone A in the treated barley (cv. Scarlett) roots was selected as a single biological replicate. The root segment was then fixed in Farmer’s fixative (EtOH: acetic acid, 3:1) under a vacuum of 500 mbar on ice for 15 minutes, followed by incubation at 4°C for 1 hour. The fixation process was repeated twice with fresh fixative solution. Subsequently, the samples were immersed in a PBS solution containing 34% sucrose and subjected to 45 minutes of vacuum treatment, followed by incubation on ice at 4°C for 21 hours. The samples were then carefully dried with tissue paper, embedded in tissue-freezing medium, and rapidly frozen in liquid nitrogen. The medium blocks containing the tissue were cut into cross sections of 20 μm thickness using a cryomicrotome (Leica CM1850). The sections were attached to PEN membrane-covered slides coated with poly-L-lysine (Zeiss). The tissue-freezing medium was removed by incubating the slides in 50% EtOH for 5 minutes, followed by sequential incubation in 70% EtOH, 95% EtOH, 100% EtOH, and 100% xylene (RNase-free) for 1 minute each. The different tissue layers were cut using the PALM Microbeam laser capture instrument (Zeiss, Germany). The cells were individually harvested and captured on the adhesive cap (Zeiss, Germany). For each tissue, more than 1,000 cells were obtained per biological replicate. RNA was isolated using the Arcturus PicoPure RNA Isolation Kit (Thermo Fisher), DNase I treatment was performed during RNA isolation using on-column DNase digestion. RNA-Seq library construction used amplification of cDNA via SMART combined with transposase-based library construction technique.

The sequencing was performed on DNBSEQ-G400 platform (BGI, China). The RNA-seq experiment produced an average of 4.42G paired-end reads per sample. The raw reads were quality checked and mapped to the barley reference genome (Barley_Morex_V2) (Mascher, 2019) by SOAPnuke (Cock *et al*., 2010) and HISAT2 (Kim *et al*., 2015). To identify the differentially expressed genes (DEGs) between different tissues, DESeq2 (Love *et al*., 2014) was used to compare gene expression levels in different tissues with FDR < 0.05 (Benjamini-Hochberg) as the criterion for DEGs.

### Weighted gene correlation network analysis

The gene coexpression network analyses were carried out using WGCNA in the R package (Langfelder & Horvath, 2008). Before the network construction, the proper soft-thresholding power was determined by an analysis of the network topology. We used the block-wise network construction option with a soft-thresholding power value of 14. To avoid small clusters, modules with < 30 genes were merged with their closest larger module using the cutreeDynamic function. The eigengene calculation of each module was performed using the moduleEigengenes function followed by the calculation of the module dissimilarity of eigengenes. Modules whose eigengenes were correlated > 0.8 were merged via the mergeCutHeight function with a cutHeight of 0.20. A unique color was then assigned to each merged module via the plotDendroAndColors function. The correlation between each gene pair was calculated to establish a similarity matrix. Using a range of soft-threshold values, the average connection network and the fitting evaluation network of scale-free topology models showed an approximate scale-free topology. The adjacency was converted into a topological overlap matrix (TOM), and all coding sequences were hierarchically clustered by TOM similarity. The Dynamic Tree Cut method, which merged highly correlated modules using a height cut of less than 0.25, was used to determine the co-expression gene modules of the gene dendrogram. Finally, we confirmed the stage-specific modules according to the distinguished interrelationship between module membership and the significance gene.

Gene-trait correlations were assessed by defining Gene Significance (GS), which quantifies the association between individual genes and external traits. Module membership (MM) was defined as the correlation between gene expression profiles and module eigengenes. Genes with the highest MM and GS values in modules of interest were identified as candidates for further investigation. Intra-modular hub genes were selected based on external traits, with criteria of GS > 0.65, MM > 0.65, and P-value < 0.05.

### Quantification of Al concentrations

Plant samples were dried at 60°C and ground into a fine powder for analysis. Using a high-accuracy balance, 100 mg of the dried and powdered plant material was transferred into a Teflon digestion tube. The samples were then digested using concentrated nitric acid (HNO₃, 65%, suprapur) and hydrogen peroxide (H₂O₂, 30%, suprapur) to ensure complete breakdown of the plant material. Trace element concentrations were determined using Inductively Coupled Plasma Mass Spectrometry (ICP-MS, iCAP TQe, ThermoScientific) and Inductively Coupled Plasma Optical Emission Spectroscopy (ICP-OES, iCAP Pro XP, ThermoScientific). To ensure the accuracy and reliability of the measurements, external standards were included alongside the samples during analysis, resulting in a relative standard deviation (RSD) of less than 2% (2σ). An internal standard (Be) was also added to each sample as a quality control measure.

### Leaf pigments and physiological parameters

Leaf pigments (chlorophyll content, flavonoid index, anthocyanin index, and nitrogen balance index) were non-destructively measured using a handheld Dualex® Scientific instrument (Force A DX16641, Paris, France). The quantum efficiencies of photosynthetic electron transport through photosystem II (PhiPS2) were measured using a portable handheld LI-600 porometer system integrated with a fluorometer (LI-COR Biosciences, Lincoln, USA). After transferring barley to the treatment solution, day 0 was set as the starting point. The measurements were conducted every day from 11:30 am to 12:00 pm. Each measurement included 9 or more biological replicates, and the same leaf position of the same leaf was used for each measurement every day.

### Statistical analysis

Statistical analyses were done with GraphPad Prism 9.0 software (https://www.graphpad.com/) or with the R Environment. For multiple comparisons between different treatment, one-way or two-way ANOVA was performed followed by Tukey’s test. Binary comparisons were performed using Student t-test.

## Results

### Effect of Al on Barley Root Suberization

To understand the impact of Al stress on suberin deposition in barley roots, we exposed 6-do barley (cv. Scarlett) plants to different conditions: a nutrient solution at pH 4.5 (control) or 5.8, and solutions with 50 μM or 100 μM AlCl_3_, for four additional days. Seminal root length remained unchanged under both pH conditions, even under acidic condition (Fig. 1A). However, Al exposure significantly reduced seminal root length (Fig. 1A).

**Fig. 1.**
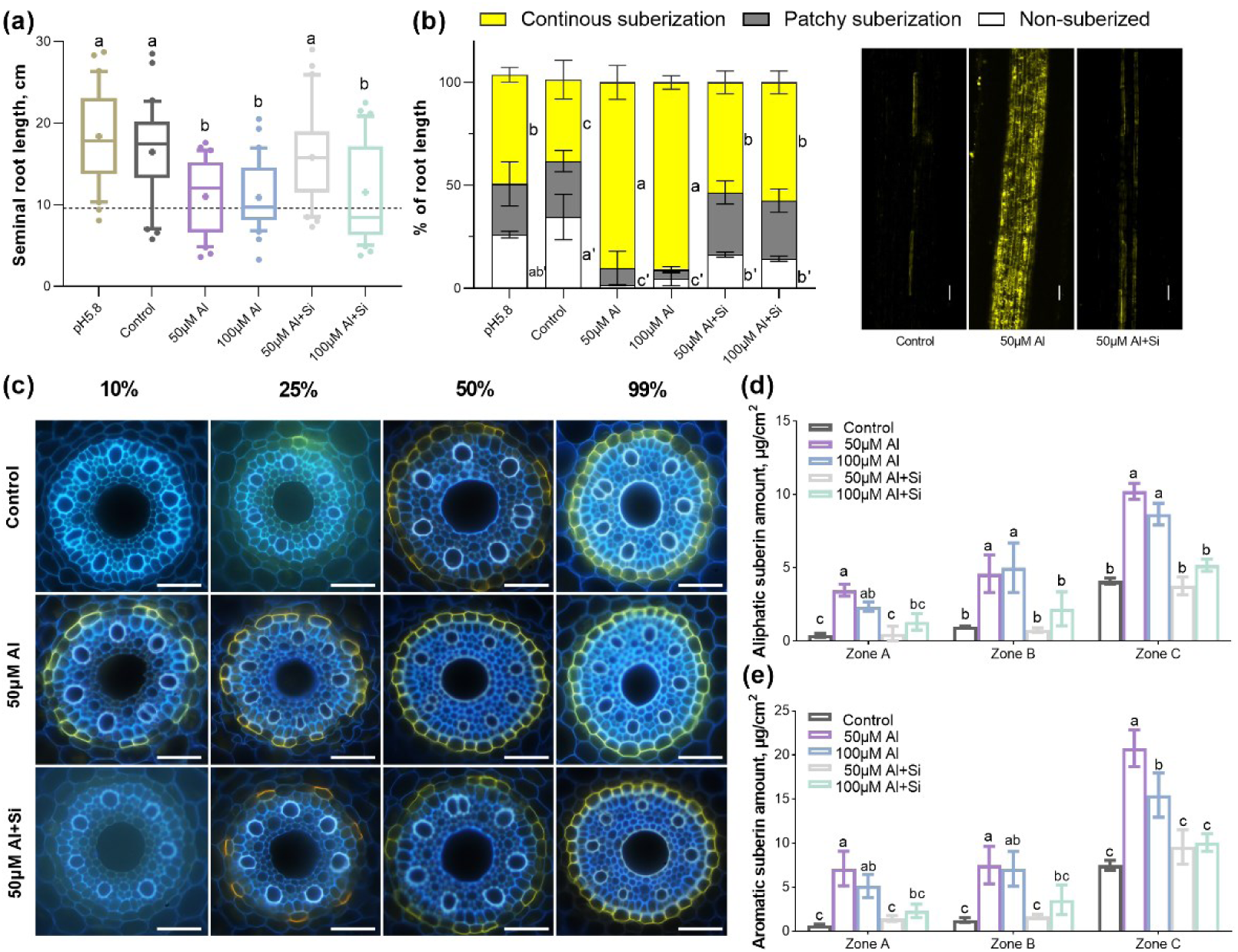
Effect of Al on root suberization of barley roots. (a) Seminal root lengths of 10-d-old barley (cv. Scarlett) plants grown under different conditions. Barley plants grown in nutrient solution at pH 4.5 were presented as controls. The boxes range from the 10 to 90 percentiles. The “+” in the box represents the mean value. The whiskers range to the outliers. Different letters indicate significant differences (P < 0.05). (b) Suberization via FY staining of barley roots under different conditions. Suberin deposition was quantified along the root axis, using three different zones: non-suberized, patchy, and continuous. Data are presented as percentages of root length. Pictures taken in similar parts of the roots. The scale bar represents 50 μm. Different letters indicate significant differences (P < 0.05). (c) Development of suberin lamellae in the endodermis of barley seminal roots. Suberin lamellae in roots grown under different conditions were stained with FY. The presence of suberin lamellae is indicated by a bright yellow fluorescence at 10%, 25%, 50%, and 99% positions of relative root length. The scale bar represents 50 μm. (d-e) Total amounts of (d) aliphatic and (e) aromatic suberin in barley seminal roots grown under different conditions. Barley plants grown in nutrient solution at pH 4.5 were presented as controls. Results are shown as mean expression ±SD of three biological replicates, different letters indicate significant differences (p< 0.05).

We used Fluorol Yellow 088 (FY) staining to detect suberin lamellae, showing bright yellow deposits in endodermal cell walls (Fig. 1B, C). Analysis indicated no suberin deposition in the youngest root zone (up to 25% of root length, Zone A), patchy suberization in Zone B (25-50%), and complete suberization in Zone C (50-100%), the mature root area, under normal conditions (Fig. 1B and Kreszies et al., 2019). Here under control conditions, barley exhibited mainly reduced and discontinuous suberization patterns (Fig. 1B). In contrast, Al-treated barley developed suberin lamellae earlier, with continuous suberization appearing at just 10% of the root length (Fig. 1B, C).

Chemical analysis after Al treatment revealed a significant increase in root suberization, consistent with microscopic observations, both aliphatic and aromatic suberin components throughout various root zones (Fig. 1D, E). In contrast, pH alterations alone did not significantly impact suberin amount in barley roots (Fig. S1, S2A). Detailed analysis of aliphatic functional groups and suberin monomers revealed marked differences, particularly elevated levels of α–ω dicarboxylic acids and ω-hydroxy acids (Fig. S2B). Further examination identified C18 α–ω dicarboxylic acids and C18 ω-hydroxy acids as the primary constituents under Al stress across all root zones (Fig. 2). The majority of ω-hydroxy acids were significantly enhanced, indicating differential regulation of suberin monomer components under Al stress, with ω-hydroxy acids particularly sensitive to Al exposure.

**Fig. 2.**
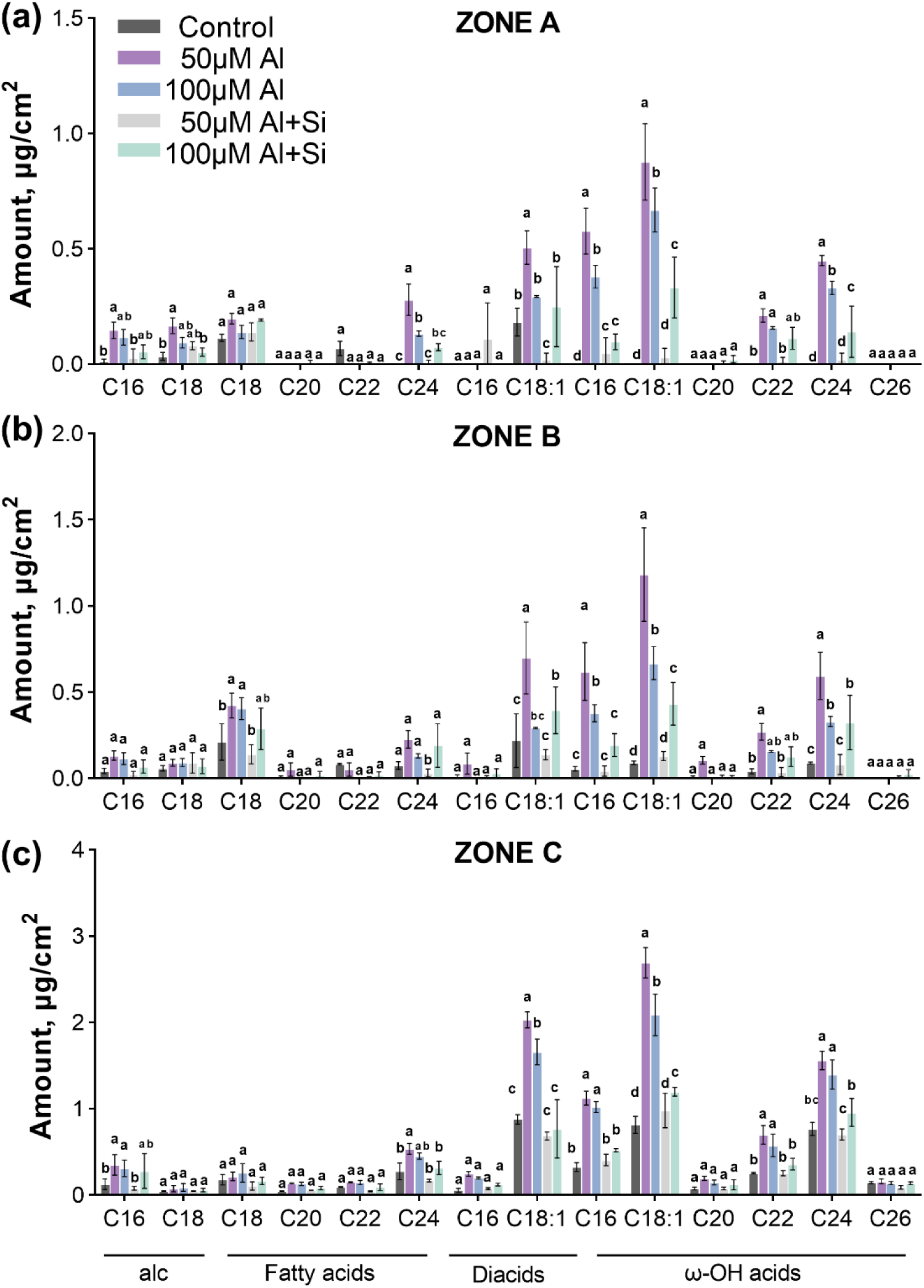
Amounts of monomers of aliphatic suberin in different zones of barley roots. Aliphatic suberin monomers amount of barley (cv. Scarlett) roots zone A (a), zone B (b), zone C (c). Plants grown under 50 μM or 100 μM Al treatment with or without Si conditions (at pH 4.5). Barley plants grown in nutrient solution at pH 4.5 presented as control. Results are shown as mean expression ±SD of three biological replicates, different letters indicate significant differences (p< 0.05). alc, primary alcohols; diacids, α–ω dicarboxylic acids; ω-OH acids, ω-hydroxy acids.

Previous studies have demonstrated that adding silicon (Si) can significantly improve plant resistance to Al toxicity (Kopittke *et al*., 2017; Hodson & Evans, 2020). In our study, the addition of Si to a medium containing 50 μM Al resulted in seminal root lengths of barley seedlings that were comparable to those in the control group. However, the growth inhibition observed with 100 μM Al was not significantly alleviated by the addition of 1 mM Si (Fig. 1A). While Si alone did not affect cell wall suberization in barley (Kreszies et al., 2020), our findings indicate that Si significantly reduced suberin deposition in the roots under Al stress (Fig. 1B, C). Moreover, Si supplementation in plants stressed with either 50 μM or 100 μM Al decreased suberin levels to those observed in the control group (Fig. 1D, E). Altogether these data demonstrate that Al stress induces root suberization in barley, by increased suberin accumulation along the root axis, rapid formation of suberin lamellae in the endodermis, and higher concentrations of both aliphatic and aromatic suberin components.

### Molecular Mechanisms of Aluminum-Induced Suberization in Barley Roots

To further explore the molecular mechanisms of root suberization induced by Al stress in barley, we collected a 5 mm segment from 25% of the root length after 4 days of Al treatment. The epidermis, cortex, endodermis, and stele cells were then isolated using laser capture microdissection (LCM), and RNA was extracted for RNA sequencing (Fig. 3A). Principal component analysis (PCA) illustrated the relationships between the transcriptomes of the Al-treated group, control group, and the four tissues (Fig. 3B). The first two components, PC1 and PC2, explained 53.1% of the total variance (Fig. 3B). The first and strongest component is due to differences between control and Al-treated samples, independently of cell type, while the second component separates the samples with a combination of treatment and cell type (Fig. 3B). Biological replicates from each treatment, including samples from the four tissues, clustered closely, indicating that transcriptomic differences among the tissues were less significant than those between the treatment and control groups.

**Fig. 3.**
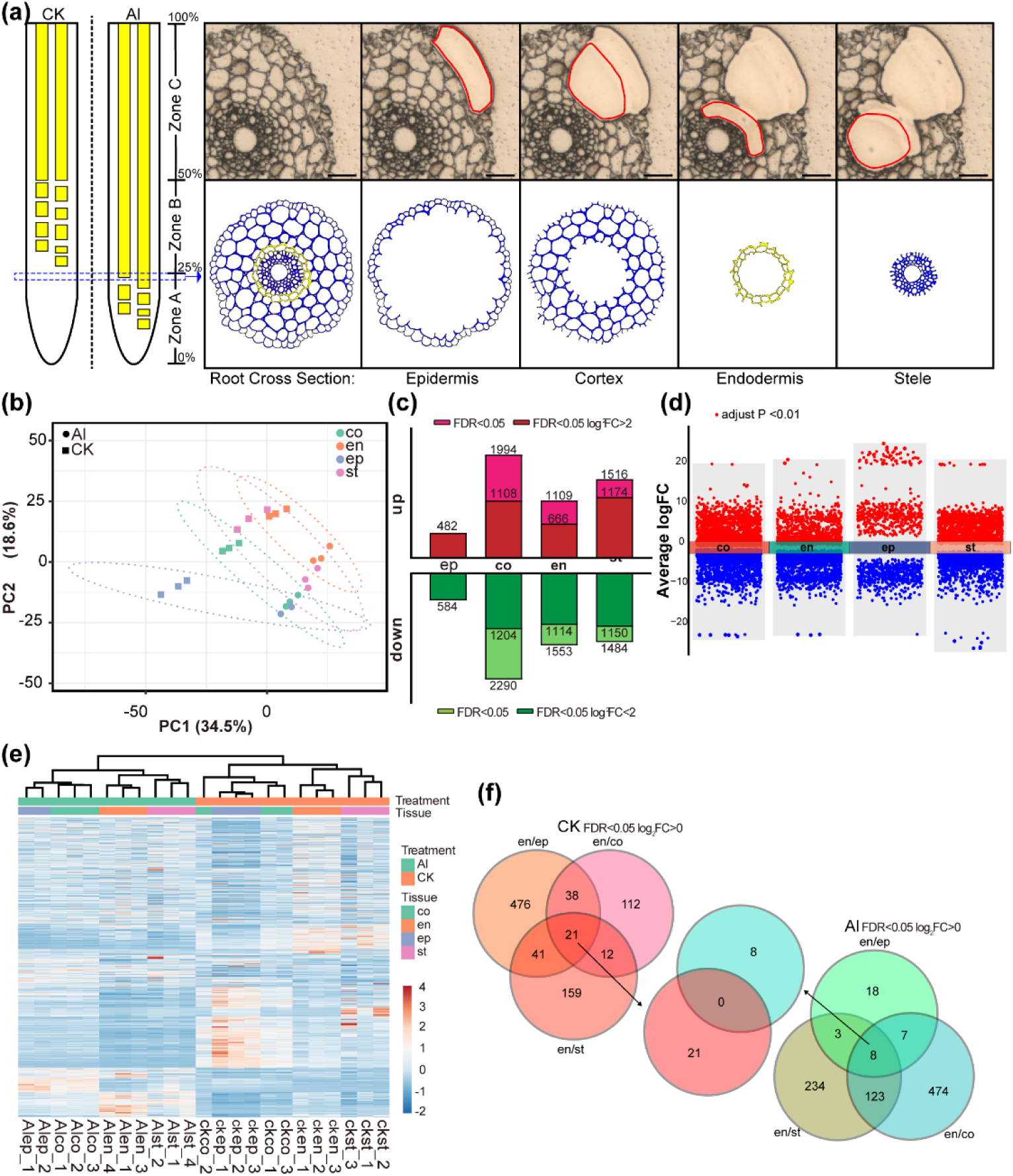
Transcriptomic analyses of the different tissues in barley seminal roots. (a) Experimental setup: RNA of root epidermis, cortex, endodermis, and stele from the 25% region of the barley (cv. Scarlett) seminal roots were isolated. The scale bar represents 50 μm. (b) PCA of the transcriptome relationships between the Al treatment group, control group, and the four tissues. PC1 and PC2 represent the two major components, explaining 53.1% of the total variance. (c) Numbers of up and downregulated genes relative to the control group under Al treatment. Light color, FDR < 0.05; Dark color, FDR <0.05, log2 fold change |log2 FC| >2. (d) Multiple volcano map of DEGs of different tissues under Al treatment relative to the control group. The Y-axis represents the log2 fold change (log2 Al /CK) of genes. Red represents DEG up-regulated, blue represents DEG down-regulated. (e) Clustering of the expression profiles of DEG between the Al treatment group, control group, and the four tissues. The color bar at the shows the normalized z scores of gene expression levels. (f) VENN diagram representing differentially expressed genes of different barley roots tissues. ep, epidermis; co, cortex; en, endodermis; st, stele.

We initially examined the expression of the *HvMATE* gene (*HORVU.MOREX.r2.2HG0137930*) and found it was strongly induced in all four tissues under Al treatment (Table S2). This response is consistent with the behaviour of Multidrug and toxin efflux (MATE) genes under Al stress observed in other species (Li *et al*., 2017), validating the effectiveness of our treatment. Differentially regulated genes (DEGs) were identified by comparing each tissue in the Al-treated group against the control, using stringent criteria (FDR < 5%, Log2FC > |0| or > |2|) (Fig. 3C and Table S2). Specifically, within the endodermis, 1109 genes were upregulated (666 with Log2FC > 2), while 1553 genes were downregulated (1114 with Log2FC < –2) (Fig. 3C-D).

The hierarchical clustering based on gene expression data revealed distinct separation between Al-treated and control samples (Fig. 3E). The cortex and epidermis grouped together, whereas the endodermis and stele formed a separate cluster. Furthermore, most genes displayed opposite expression trends under Al treatment compared to the control group (Fig. 3E).

To further pinpoint endodermis-specific genes responsive to Al stress, we conducted a comparative analysis of DEGs across various tissues (Fig. 3F). We identified 36 genes with an endodermis/epidermis Log2FC > 0 (FDR < 0.05), 612 genes with an endodermis/cortex Log2FC > 0 (FDR < 0.05), and 368 genes with an endodermis/stele Log2FC > 0 (FDR < 0.05). The overlap was 8 genes, which were identified as endodermis-specific genes under Al treatment. In contrast, under control conditions, 21 genes were identified as endodermis specific. Intriguingly, there was no overlap between the genes upregulated under Al stress and those in control conditions (Table. S3).

Kyoto Encyclopedia of Genes and Genomes (KEGG) enrichment analysis of endodermis DEGs highlighted 17 genes associated with cutin, wax, and suberin synthesis pathways (Fig. 4A, Table S4), including seven Cytochrome P450 genes (Fig. 4B). A phylogenetic analysis identified *HORVU.MOREX.r2.1HG0034810* (*HvCYP86B1*) as the barley homolog of the Arabidopsis *CYP86B1* gene. Notably, *HvCYP86B1* expression was significantly higher in the endodermis and cortex of the control group compared to the Al-treated group (Fig. 4C). We also mined publicly available transcriptomes in barley treated with Al or low pH (Szurman-Zubrzycka *et al*., 2021). Several key suberin-biosynthesis genes have been shown affected by Al treatment. Under long-term (7 days) Al exposure, suberin-biosynthesis genes such as *CYP86B1*, *KCS1*, and *GPAT4* were notably upregulated (Fig. S3). In contrast, no significant changes were observed under low pH or short-term (24h) Al treatments, indicating differential gene regulation in response to prolonged Al exposure.

**Fig. 4.**
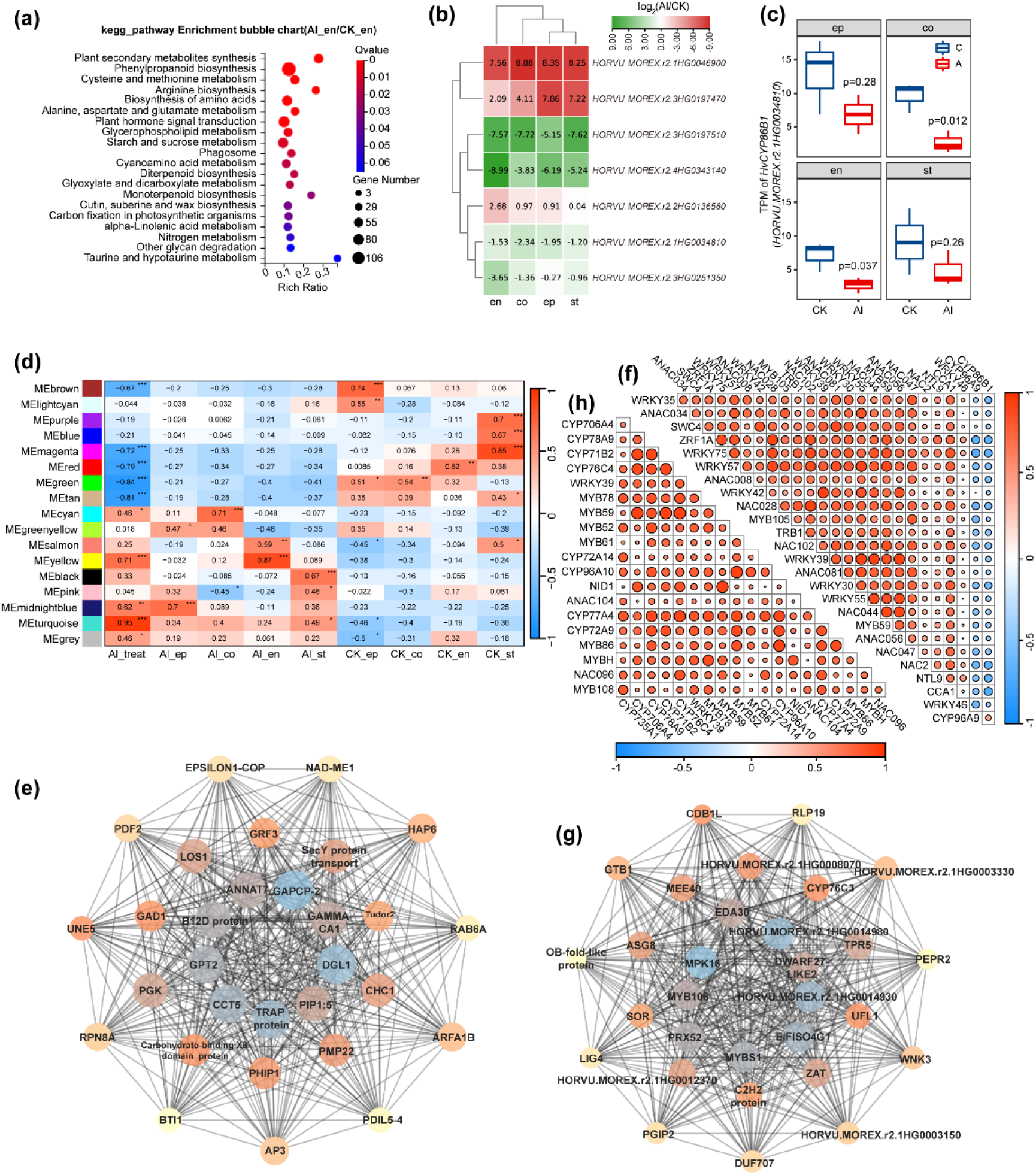
Tissue-specific transcriptomic responses and regulatory networks involved in suberin biosynthesis under Al stress. (a) KEGG enrichment analysis of DEGs in the endodermis under Al treatment relative to the control group. (b) Heatmap of log_2_ fold change for seven Cytochrome P450 family protein genes in four different tissues. (c) TPM of *HvCYP86B1* in four different tissues under Al treatment and control. ep, epidermis; co, cortex; en, endodermis; st, stele. (d) Module–trait relationships. The Al treatment (Al_treat) and four tissues (ep, epidermis; co, cortex; en, endodermis; st, stele) are used as traits, each column corresponds to a different trait. Each row corresponds to the characteristic genes of the module. The relationship between the modules and traits is indicated in cell by Pearson’s correlation coefficients. Asterisks indicate significant values calculated using the corPvalueStudent function: *, *P* < 0.05; **, *P* < 0.01; ***, *P* < 0.001. Cell color ranges from red (highly positive correlation) to blue (highly negative correlation). (e) co-expression network of top30 hubgenes in module Meturquoise. (f) Pearson correlation analysis between suberin biosynthesis-related genes and transcription factors identified in the hub gene modules under Al stress. The heatmap displays the Pearson correlation coefficients, where red indicates a positive correlation and blue indicates a negative correlation. The intensity of the color corresponds to the strength of the correlation. (g) co-expression network of top30 hubgenes in module Meyellow. (h) Pearson correlation analysis between suberin biosynthesis-related genes and transcription factors identified in the hub gene modules under Al stress in endodermis.

To validate these findings, we examined the expression of suberin-related genes in barley roots subjected to different treatments using RT-qPCR (Fig. S4). We initially analyzed two key suberin synthesis genes, *HORVU.MOREX.r2.3HG0251350* (*HvCYP86A1*) and *HvCYP86B1*, under Al treatment with or without Si added in different root zones. Under Al treatment, *HvCYP86B1* was upregulated in zone A and B, while *HvCYP86A1* was upregulated only in zone A, with no significant changes in other zones, consistent with previous transcriptomic sequencing results.

To reveal additional genes that are significantly associated with *Al treatment* and different tissuses of barley seminal roots, we performed a weighted gene co-expression network analysis (WGCNA). This approach identified 17 co-expression modules within the root, among which the MEgrey module contained transcripts that did not meet the selection criteria (Fig. 4D). These modules were analyzed for their association with Al treatment and specific root tissues. Specifically, MEturquoise module exhibited the highest positive correlation with Al treatment, marking it as the key module associated with Al-responsive genes. The MEyellow module demonstrated the strongest correlation with both Al treatment and the endodermis, making it the primary module for analyzing Al-induced gene expression in the endodermis.

In the MEturquoise module, Module Membership (MM) and Gene Significance (GS) were positively correlated (Fig. S5A), indicating that the genes in this module are highly associated with the Al treatment trait and play critical roles in Al-responsive mechanisms. The module contained 6,303 genes, and by applying thresholds of GS > 0.65 and MM > 0.65, we identified 2,896 hub genes (Table. S5), including *HvCYP86B1* and *HORVU.MOREX.r2.4HG0343140* (*HvCYP96A9*), both of which are linked to suberin biosynthesis. Co-expression network analysis of the top 30 hub genes revealed their central roles in the response to Al stress (Fig. 4E). By predicting transcription factors for these 2,896 hub genes using protein sequences from the PlantTFDB database, we identified 94 transcription factors, with 24 belonging to the MYB, WRKY, and NAC families (Table. S5). Pearson correlation analysis was then performed to explore the relationships between *HvCYP86B1*, *HvCYP96A9*, and the transcription factors identified within the hub genes (Fig. 4F). The results revealed that *HvCYP86B1* and *HvCYP96A9*, key genes involved in suberin biosynthesis, were predominantly negatively correlated with MYB, WRKY, and NAC transcription factors. This suggests that these transcription factors may act as negative regulators of suberin biosynthesis in barley roots under Al stress.

In the MEyellow module, MM and GS were also positively correlated (Fig. S5B), suggesting a strong association between this module’s genes and the Al-endodermis trait. The MEyellow module contained 1,057 genes, including nine cytochrome P450 family members such as *HORVU.MOREX.r2.2HG0136560* and *HORVU.MOREX.r2.3HG0197470*. Transcription factor prediction identified 28 transcription factors in the module, 11 of which belonged to the MYB, WRKY, and NAC families (Table. S5). Co-expression network analysis of the top 30 hub genes indicated that *HvMYB108* was a key hub gene in the MEyellow module (Fig. 4G).

Further correlation analysis of the nine cytochrome P450 genes and the transcription factors identified among the hub genes revealed significant positive correlations between potential suberin biosynthesis genes and transcription factors from the MYB, WRKY, and NAC families, particularly *MYB108*, *MYB86*, *MYB61*, *MYB52*, *MYB59*, and *MYB78* (Fig. 4H). In addition, the nine cytochrome P450 genes identified as potential regulators of suberin biosynthesis provide new candidates for future experimental studies of suberin pathways in barley.

Collectively, these findings illustrate that Al stress induces distinct molecular responses in barley roots, leading to increased suberin deposition. This process is characterized by the upregulation of key suberin biosynthesis genes, particularly under prolonged Al exposure.

### Impact of Al on Suberin Development in Barley Loss-of-Function Mutants

To further understand the impact of Al on suberin development, we employed two barley loss-of-function mutants, *cyp86b1-1* and *cyp86b1-2*, generated in the Golden Promise Fast (GPF) cultivar. These mutants have a 170 bp deletion and a 168 bp inversion, respectively. The gene encoded by *cyp86b1-1* produces only 107 normal amino acids (out of 548) because the deletion not only removes the sequence coding for amino acids 108 to 164 but also changes the reading frame to produce a completely different protein sequence. In *cyp86b1-2*, the inverted region changes the putative protein sequence after amino acid 107 and it truncates it after 31 amino acids because of an early stop codon. (Fig. 5A, S6). Thus, it is expected that both alleles result in complete loss of HvCYP86B1 activity. We compared the growth of these mutants to GPF under both Al and control conditions. There was no significant difference in root or shoot length between the genotypes under control conditions (Fig. 5B, C). Moreover, Al treatment significantly inhibited root growth (length and biomass) in both mutants and in GPF plants, but the effect was stronger in the mutants (Fig. 5B, S7A). On the other hand, Al treatment did not affect shoot length nor weight in any of the genotypes (Fig. 5C, S7B). We also analyzed the suberin amount in the roots. Notably, the suberin composition and amounts in GPF control were similar to those in Scarlett (Fig. 1D, E, Fig. 5D, and Fig. S8). When comparing the GPF with the mutants under control conditions, there was no significant difference in total aliphatic suberin amount between the genotypes (Fig. 5D). However, a detailed analysis of aliphatic suberin monomers revealed that in the mutants, C22 and C24 ω –hydroxy acids were almost absent, but fatty acids accumulated in greater amounts (Fig. S9). After Al treatment, the aliphatic suberin amount in all three zones of the *cyp86b1-2* mutant roots was lower than in GPF, while in the *cyp86b1-1* mutant, only the aliphatic suberin amount in zone C was lower than in GPF (Fig. 5D). Similar to the control conditions, the C22 and C24 ω-hydroxy acids in the mutants were almost absent, accompanied by fatty acid accumulation (Fig. S10). Among the four aliphatic suberin components, there was no significant difference in amount between GPF and mutant barley in the control group, except for a lower ω-hydroxy amount in the mutants in zone C (Fig. S11). The primary alcohol amount remained unchanged under Al treatment in both GPF and mutant barley, while the fatty acids in zones A and B were lower in GPF compared to the mutants. Conversely, for α-ω dicarboxylic acids and ω-hydroxy acids, the contents were significantly lower in the mutants.

**Fig. 5.**
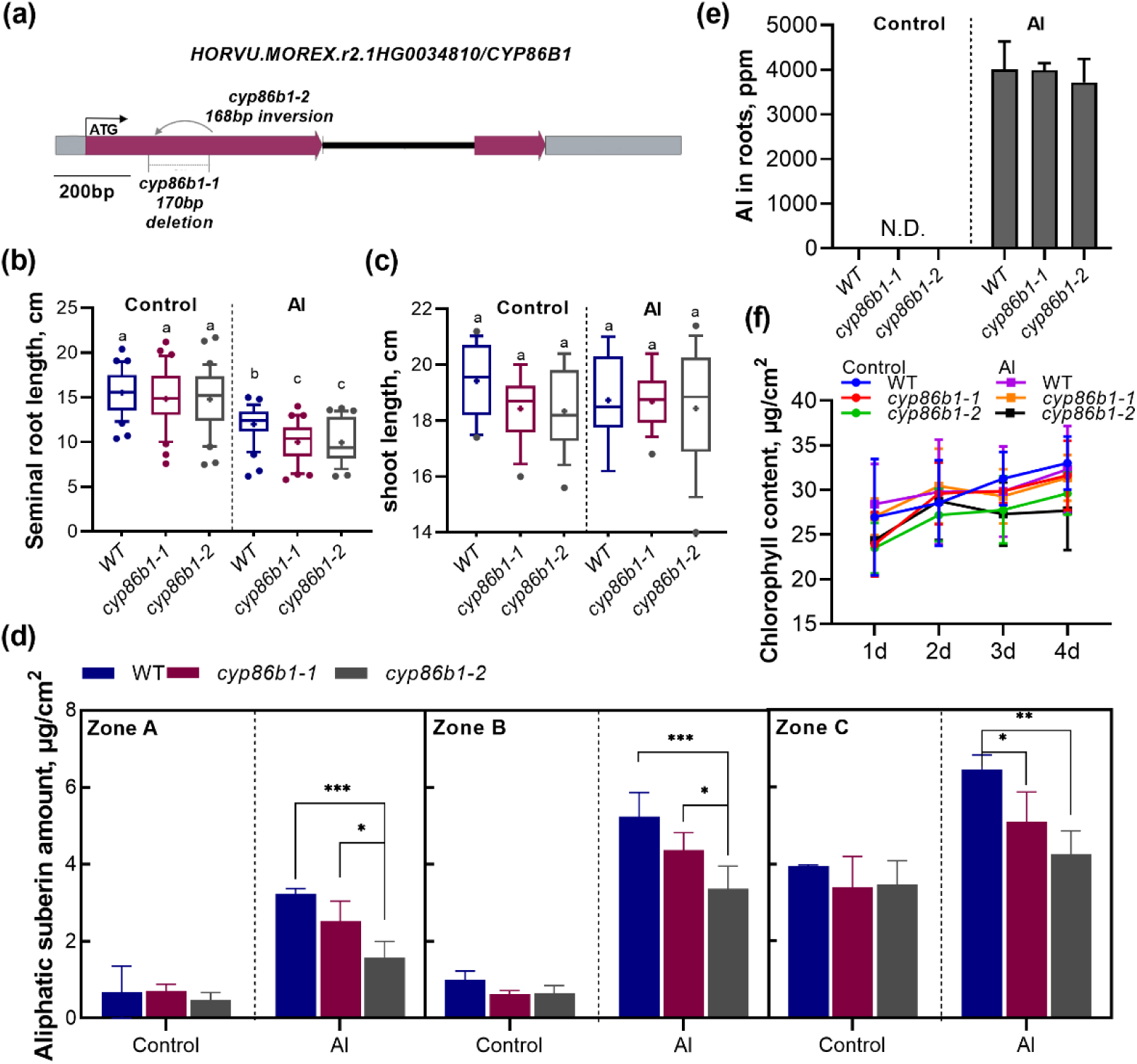
Impact of Golden Promise Fast CYP86B1 mutations. (a) Gene structure of CYP86B1 (HORVU.MOREX.r2.1HG0034810) with mutations in cyp86b1 (cyp86b1-1: deletion and cyp86b1-2: inversion). Seminal root lengths (b) and shoot lengths (c) of barley plants grown in nutrient solution (control) or containing 50 μM AlCl_3_ (Al) at pH 4.5 for 4 days. The boxes range from the 10 to 90 percentiles. The “+” in the box represents the mean value. The whiskers range to the outliers. Error bars represent SD, different letter indicate significant differences (p< 0.05). (d) Total amounts of suberin aliphatic in GPF and barley mutant roots. Barley plants were grown in nutrient solution (control) or containing 50 μM AlCl_3_ (Al) at pH 4.5 for 4 days. Results are shown as mean expression ±SD of three biological replicates, *P < 0.05; **P < 0.01; ***P < 0.001. (e) Al concentration in GPF and barley mutant shoots under Al treatment. Results are shown as mean expression ±SD. “N.D.” indicates that no detectable levels were found. No significant differences were detected between mutants and GPF plants under each condition. (f) Effect of Al stress on chlorophyll content in GPF and barley mutant shoots. Results are shown as mean expression ±SD. No significant difference was detected.

We then analyzed the impact of reduced Al-induced suberin in *cyp86b1* mutants on barley’s absorption of Al. First, we examined the Al concentration in barley roots exposed to Al and found no difference between the genotypes (Fig. 5E). Then, to investigate the effects of Al and suberin on barley shoot development, we analyzed leaf chlorophyll content and other leaf physiological indicators but found no differences between the genotypes (Fig. 5F, Fig. S12). Altogether, the analyses suggest that reduced suberin accumulation in *cyp86b1* mutants under Al stress, does not affect Al absorption or pigment content in barley.

### Al-induced suberization is influenced by the ABA Pathway

As suberin is induced by ABA and SGN3/CIFs independently (Shukla *et al*., 2021), we tested which of these pathways mediates Al-induced changes in suberization. We did not observe any differential expression of barley SGN3 homologous genes in our LCM RNA-seq results with barley. However, our analysis of LCM RNA-seq revealed significant changes in ABA-related genes due to Al treatment (Table S2 and Fig. S13). This suggests that ABA signalling plays a role in barley’s response to Al stress. Similarly, qPCR results showed a broad response of ABA pathway genes under Al treatment in different barley (cv. Scarlett) root zones (Fig. 6A).

**Fig. 6.**
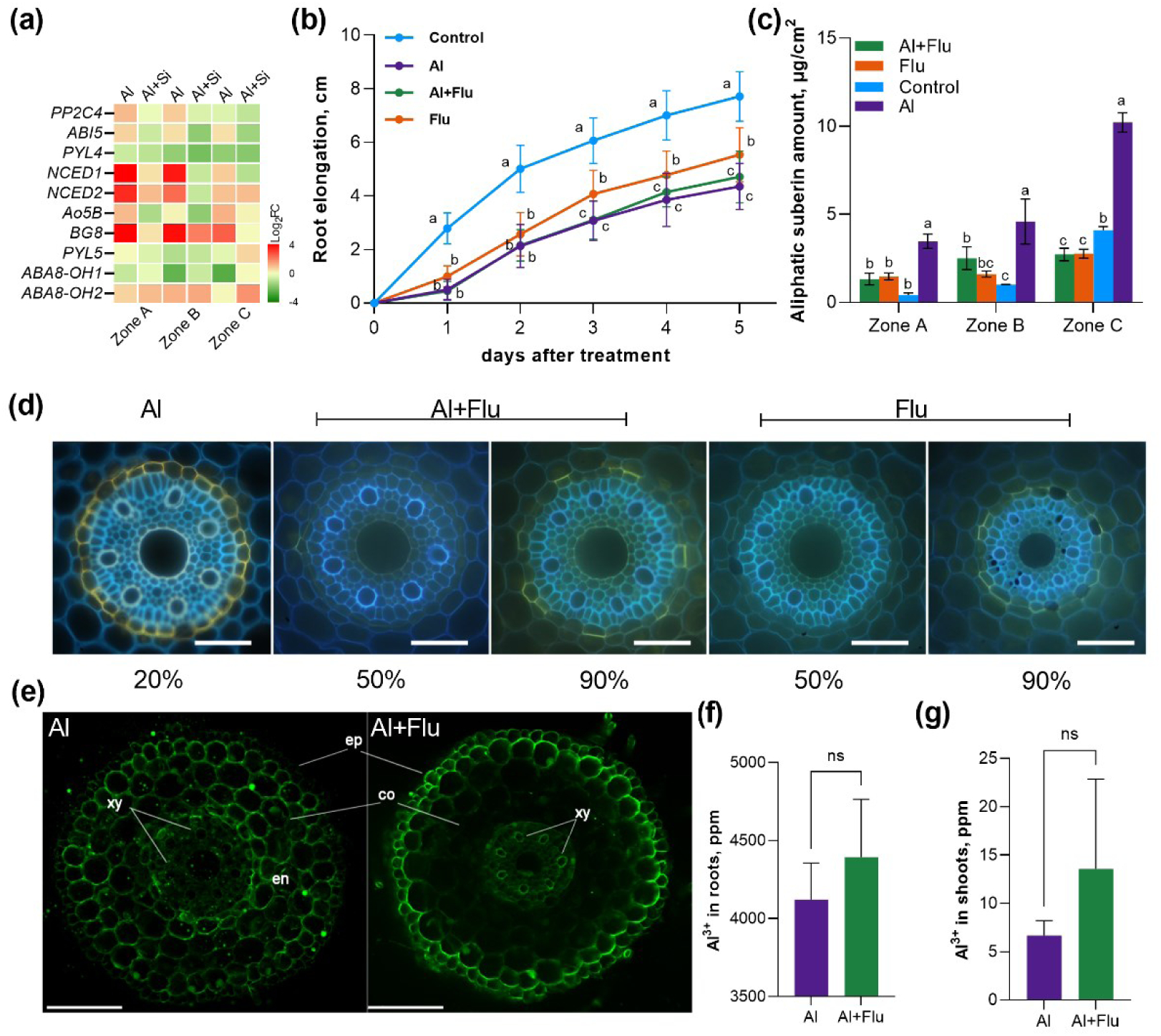
Effect of suberin on Al uptake in barley seminal roots (cv. Scarlett). (a) Relative expression levels of the ABA-related genes in barley roots. Barley roots were treated with 50 μM AlCl_3_ or 50 μM AlCl_3_ with 1mM Si additions for 4 days. Results are presented as Log2 fold changes compared with the control condition. (b) Root growth of barley plants grown under different treatment conditions. Results are shown as mean expression ±SD (n ≥ 50 seminal roots), different letters indicate significant differences (p< 0.05). (c) Total amounts of aliphatic suberin in barley roots grown under different treatment conditions. Results are shown as mean expression ±SD of three biological replicates, different letters indicate significant differences (p< 0.05). (d) Development of suberin lamellae in the endodermis of barley seminal roots. The presence of suberin lamellae is indicated by a bright yellow fluorescence. Blue fluorescence indicates autofluorescence. Pictures were taken at 20%, 50% or 90% of relative root length. The scale bar represents 50 μm. (e) Subcellular distribution of Al. Al distribution in roots grown under different conditions was stained with morin (green fluorescence). Roots were sectioned at 20% of root length from the apex for morin staining and fluorescence observation. Bar = 100 µm. ep, epidermis; co, cortex; en, endodermis; xy, xylem. Al concentration in barley roots (f) and shoots (g) under Al treatment with or without 10 μM Flu. Results are shown as mean expression ±SD. No significant difference was detected.

To further explore the role of ABA in Al-induced suberin formation, we examined the impact of fluridone (Flu), an inhibitor of ABA biosynthesis in barley and other gramineous plants (Gamble & Mullet, 1986). Compared to the control group, Flu treatment significantly inhibited barley (cv. Scarlett) root growth (Fig. 6B). Al exposure inhibits root growth even more strongly than Flu but the combination of Al and Flu did not impair root length any further (Fig. 6B). Next, we analyzed the effects of Flu on suberin biosynthesis under Al treatment. The suberin amount in barley roots showed no significant difference between the Flu alone treatment and the Flu combined with Al treatment. However, the suberin levels after Flu combined with Al treatment were lower than those observed with Al treatment alone (Fig. 6C, S14). Additionally, FY staining of barley seminal root cross-sections showed that Flu treatment significantly delayed the appearance of the suberization zone, with the first occurrence of single suberized cells only within the range of about 50-60% of root length (Fig. 6D). The pattern of suberization formation under Al treatment combined with Flu was similar to that observed with Flu treatment alone. Thus, Flu significantly inhibited the promoting effect of Al on suberization.

To explore the role of suberin in Al transport within barley roots, we used morin staining to detect Al entry on cross-sections of root apices (Zone A). Green fluorescence, indicative of Al presence, was predominantly observed in the root cortex, specifically external to the endodermis, after 4 days of Al treatment (Fig. 6E). However, in samples co-treated with Flu and Al, green fluorescence in the cortex was notably reduced, although it increased within the stele, particularly the xylem, and in the outermost layers of the root.

Following the morin staining results, Al concentration in barley roots was analyzed. The results indicated no statistically significant difference in Al concentration between roots treated with Al and Al+Flu. Similarly, analysis of the aboveground parts of barley revealed a comparable trend: there was no statistically significant difference in Al concentration between the shoots and roots under Al+Flu treatment and Al treatment alone.

## Discussion

Barley is one of the most Al-sensitive cereal species, with notable variation in Al tolerance among different cultivars. This tolerance largely correlates with the ability of the genotype to secrete citrate (Zhao *et al*., 2003; Furukawa *et al*., 2007). The cultivar ‘Scarlett’ used in our study is relatively sensitive to Al. Even micromolar concentrations of Al drastically reduce total root length (Vega *et al*., 2019). However, the reduction in root growth is not only caused by Al toxicity but also by too low pH and H^+^ toxicity, as barley is known to be very sensitive to H+ toxicity as well (Zhao *et al*., 2003; Guo *et al*., 2004). Our results show that barley seedlings tolerate a moderate decrease in pH from 5.8 to 4.5 without significant effects on root development, indicating a high buffering capacity and adaptability to acidity changes. However, the addition of Al led to a marked decrease in root length and dry weight, suggesting that Al toxicity is the primary factor limiting root growth under acidic conditions. This reduction can be attributed to inhibited cell division and elongation, and disrupted nutrient uptake and transport (Kochian *et al*., 2015). Hence, Al stress proves more detrimental to barley root development than pH stress alone. The relationship between root suberization and pH is not well understood, but some studies have suggested that suberin deposition may affect or be affected by pH in different ways (Feng *et al*., 2022). For instance, it has been suggested that low pH inhibits some crucial enzymes involved in suberin synthesis and polymerization (van Doom & Perik, 1990). Our study found that suberin lamellae distribution and amount in barley (cv. Scarlett) roots did not significantly differ between pH 4.5 and pH 5.8 after four days of treatment (Fig. S1, S2A). However, it might be that longer acidification periods may further inhibit suberin synthesis genes.

Plants can adjust suberization levels in response to various nutritional stresses. Excess salt, drought, and deficiencies in elements like K, Fe, or S can alter endodermal suberization (Barberon *et al*., 2016). Our study found that Al strongly induces suberization (Fig. 1, 2), with upregulation of suberin synthesis genes occurring after prolonged Al stress. In our sampled sites, roots treated with Al exhibited fully formed suberin lamellae, whereas the control group showed cells on the verge of suberization. We propose that this is the cause of lower expression of suberin synthesis-related genes in the Al-treated group compared to the control in our RNA-seq data, which highlights the temporal lag between suberization and corresponding gene expression changes. Conducting LCM RNA-seq analysis on samples closer to the root tip may provide a more direct correlation between suberin synthesis genes and Al treatment. Suberization forms a protective barrier in the endodermis, limiting water and solute transport in root tissues (Franke & Schreiber, 2007). The formation of suberin lamellae in barley roots was accelerated and intensified under Al stress conditions, especially at higher concentrations of Al. This suggests that suberization is an effective response to Al stress in barley, which may help to reduce the entry and accumulation of Al in the root tissues. However enhanced suberization and delayed root growth in response to abiotic stresses might also negatively affect water and nutrient uptake (Ranathunge *et al*., 2011). Therefore, suberization may be a trade-off between protection and performance in barley roots under Al stress. Nevertheless, under Al stress it seems a good strategy for the plant to investigate the metabolic costs of building up the suberin polymer in order to survive.

The aliphatic suberin monomers in barley are mainly composed of α–ω dicarboxylic acids, ω-hydroxy acids, primary alcohols, and fatty acids, derived from fatty acid metabolism (Kreszies *et al*., 2019). Al stress induced a significant increase in α–ω dicarboxylic acids and ω-hydroxy acids, especially in the youngest root zone, suggesting enhanced biosynthesis of the suberin polyester backbone (Fig. 2, S2B). This increase, particularly in the predominant C18 and C16 chain lengths, likely confers higher rigidity and stability to suberin, enhancing its protective role against Al stress (Graça & Pereira, 2000; Pollard *et al*., 2008).

Si, though not essential but beneficial, plays a crucial role in reducing Al toxicity in plants (Coskun *et al*., 2019). Our study showed that Si application alleviated Al-induced suberization in barley roots. Moreover, while the addition of 1 mM Si restored the growth of roots treated with 50 μM Al to normal levels, no restoration occurred at 100 μM Al. However, Si supplementation at both 50 μM and 100 μM Al stress reduced suberin levels in roots back to those observed in the control group. This suggests that Si might mitigate Al-induced root growth inhibition and suberization through different mechanisms. These may involve the formation of Al-Si complexes in root cell walls or apoplasts, reducing free Al concentration and preventing its uptake by root cells (Hodson & Evans, 2020). This reduced Al uptake may then decrease the need for suberization as a defense mechanism, resulting in lower suberin content in barley roots treated with Si. When Si is added, the formed Si crystals might take over some of the barrier function of suberin, potentially saving metabolic costs. Alternatively, Si may modulate the expression of genes involved in suberin biosynthesis or degradation (Fleck et al., 2011; Hinrichs et al., 2017; Vulavala et al., 2016; Wu et al., 2019), affecting suberin accumulation in barley roots under Al stress. Further studies are needed to elucidate the molecular mechanisms of Si-mediated alleviation of suberization in barley roots under Al stress.

By employing LCM followed by RNA-seq, we were able to isolate and examine the molecular responses in distinct root tissues, including the epidermis, cortex, endodermis, and stele. This approach provided high resolution in uncovering tissue-specific gene expression changes under Al stress. Our findings demonstrate that Al stress induces significant transcriptomic reprogramming in all root tissues, with pronounced differences in gene expression between Al-treated and control samples (Fig. 3). Notably, the endodermis showed substantial changes in gene expression, including a strong induction of genes associated with suberin biosynthesis. Through comparative analysis, we identified eight endodermis-specific genes responsive to Al stress, none of which overlapped with genes upregulated under control conditions, suggesting that these genes play a specialized role in the Al-induced defense mechanism (Fig. 3F).

We identified several cytochrome P450 family genes, including *HvCYP86B1* and *HvCYP96A9*, as key regulators of suberin biosynthesis under Al stress. A substantial number of genes related to suberin biosynthesis were significantly upregulated in response to prolonged Al exposure, while shorter exposure durations or low pH conditions did not elicit the same response (Fig. S3). This highlights the specific and prolonged nature of the Al-induced regulatory pathways. Additionally, the identification of MYB, WRKY, and NAC transcription factors that are negatively correlated with suberin biosynthesis genes in the MEturquoise module suggests a potential regulatory network controlling suberin deposition in response to Al stress. Conversely, in the MEyellow module, MYB transcription factors, including hub gene *HvMYB108* showed positive correlations with suberin biosynthesis genes. Therefore, the MYB108 could be key targets for further research into the molecular mechanisms of suberin biosynthesis under Al stress in barley.

The *CYP86B1* gene, encoding a cytochrome P450 monooxygenase, is involved in suberin biosynthesis (Compagnon *et al*., 2009). Among the aliphatic suberin components, the ω-OH and diacids with chain lengths higher than C18, are considered to be the major products of *CYP86B1*-mediated hydroxylation. Therefore, the lower levels of ω-OH and diacids in the *cyp86b1* mutants indicate that the CYP86B1 enzyme is essential for their biosynthesis. Moreover, the absence of C20-C26 ω-OH in the mutants verifies that CYP86B1 is responsible for the formation of ω-OH beyond C18. Although the long-chain compounds are affected, the total amount of aliphatic suberin is not affected because the shorter compounds compensate for the loss of the longer ones. The lack of C20-26 ω-hydroxy acid synthesis in *cyp86b1* mutants leads to the accumulation of precursor substance, C20-24 FA (Fig. S9, S10), which is not sufficient to protect the roots from Al stress, as evidenced by the lower aliphatic suberin amount in the mutants under Al treatment (Fig. 5).

Al stress did not affect photosynthesis and pigment metabolism in barley shoots in this study, contrasting with previous reports. The discrepancy may be due to the mild Al concentration used (Panda *et al*., 2003) and the young plant age in our study. Also, most Al is bound to root cell walls and is not very mobile within the plant, which leads to 300 to 800 times lower Al concentration in the barley shoots compared to the roots (Fig 5F, G). The suberin-deficient mutants showed no difference in shoot performance compared to GPF plants, suggesting suberin’s role in shoot protection is limited under Al stress. However, this may also be due to the relatively high suberin content still present in the *cyp86b1* mutants. Nonetheless, suberin might subtly influence shoot physiology and metabolism, for example through hormonal balance and antioxidant defense.

Flu treatment alone significantly inhibited root elongation, since ABA is essential for normal root development. However, Flu did not mitigate Al’s inhibitory effect on root growth, indicating that ABA is not involved in Al-induced growth inhibition (Fig. 6B). Previous studies link ABA to enhanced suberization (Barberon *et al*., 2016), and our results show that many ABA-related genes are regulated under Al stress (Fig. 6A). However, the SGN3/CIFs pathway, another regulator of suberin development (Shukla *et al*., 2021), did not exhibit changes under Al stress, leading us to conclude that Al-induced suberization is controlled independently by the ABA pathway.

Flu treatment reduced suberin amount and delayed suberization in barley roots under Al stress. Analysis of Al concentration in barley roots indicated that roots treated with both Al and Flu had slightly higher Al concentrations compared to those treated with Al alone, although this difference was not statistically significant. Similarly, in the aboveground parts of barley, the shoots accumulated more Al ions under the Al+Flu treatment compared to Al treatment alone, but again, these differences were not significant. Although no significant differences in Al concentrations were detected between roots treated with Flu and those treated with both Flu and Al (Fig. 6F), the analyses were limited to whole roots. Morin staining, which specifically targets Zone A, did not enable quantification of Al concentrations in this specific zone. However, it was revealed that Flu significantly affected Al distribution and transport in barley roots under Al stress, reducing Al accumulation in the root cortex but increasing it in the central cylinder. This suggests that suberin acts as a barrier, limiting Al entry and transport in the roots. These findings suggest that suberin may influence the distribution and accumulation of Al within different parts of the barley plant, potentially altering its translocation from roots to shoots. Collectively, these results highlight the role of the ABA pathway in modulating suberin synthesis under Al stress in barley and indicate that suberin impacts Al distribution within the plant, affecting its overall tolerance to Al stress.

Suberin may also bind metal ions and influence transporters involved in Al uptake and efflux (Krishnamurthy *et al*., 2009; Machado *et al*., 2013; Liska *et al*., 2016). We also observed morin stain in the outer layers of roots treated with Flu+Al, but no exodermis formation was observed under this treatment. The reasons behind this altered Al accumulation pattern remain unclear. We speculate that Flu treatment may induce changes in the expression of Al transporter proteins or mate transporters at the epidermis, leading to Al deposition in this region. These findings emphasize that our understanding of the root physiological mechanisms of Al are still poorly understood and that further research is needed to comprehend the mechanisms behind suberin-mediated Al resistance. In summary, our findings highlight the complex interplay between suberization, Al stress, and Si application in barley, offering insights into potential strategies for improving crop resilience to Al toxicity.

## Acknowledgement

The authors are grateful to Dr Li Guo and Professor Frank Hochholdinger (Institute of Crop Science and Resource Conservation, University of Bonn, Bonn, Germany) for assisting in performing LCM and to Edelgard Wendeler for technical support with barley transformation. This work was funded by the Deutsche Forschungsgemeinschaft (DFG, German Research Foundation) PN511193270 and INST 217/939-1 FUGG. Financial support from the China Scholarship Council (CSC), is highly appreciated.

## Data availability

All study data are included in the article and/or supplementary Material. The raw sequencing data have been deposited at the National Center for Biotechnology Information (NCBI) sequence read archive (SRA accession: PRJNA992920).

## Author contributions

HM and LS planned and designed the research; HM, QZ, and TK performed the research; IA produced genetic materials; HM, QZ, and TK analyzed the data; and HM and LS wrote this paper. All authors revised the manuscript and approved the final manuscript.

## Supplementary data

**Table S1.** Primers used in this study.

**Table S2.** Complete list of differentially expressed genes.

**Table S3.** Epidermis specific genes.

**Table S4.** KEGG enrichment analysis of cutin suberin wax pathway in the barley endodermis in response to Al stress.

**Table S5.** WGCNA data.

**Table S6.** DEGs and TPM values of barleys in response to Al stress.

**Fig. S1.** Total amounts of suberin in barley roots grown under different pH conditions.

**Fig. S2.** Amounts of substance classes of aliphatic suberin in barley seminal roots grown under different conditions.

**Fig. S3.** Expressions of selected suberin-biosynthesis genes of barley under different conditions.

**Fig. S4.** Expression analysis of HvCYP86A1 and HvCYP86B1 genes of barley using qPCR.

**Fig. S5.** The correlation between Module Membership and Gene Significance.

**Fig. S6.** Protein sequence of *cyp86b1* mutants.

**Fig. S7.** Effect of Al on shoot and root dry weight of wild type and transgenic barley plants.

**Fig. S8.** Aromatic suberin amount in barley mutant.

**Fig. S9.** Amounts of monomers of aliphatic suberin in different zones of barley mutant roots.

**Fig. S10.** Amounts of monomers of aliphatic suberin in different zones of barley mutant roots under Al conditions.

**Fig. S11.** Amounts of substance classes of aliphatic suberin in wild type and transgenic barley seminal roots.

**Fig. S12.** Effect of Al stress on Flavonoid index, Anthocyanin index, Nitrogen Balance Index of GPF and barley mutants.

**Fig. S13.** Expressions of selected ABA pathway genes of barley under different conditions.

**Fig. S14.** Aromatic suberin amount in barley roots under Flu treatment.

---

**Supplemental Fig. S1.**
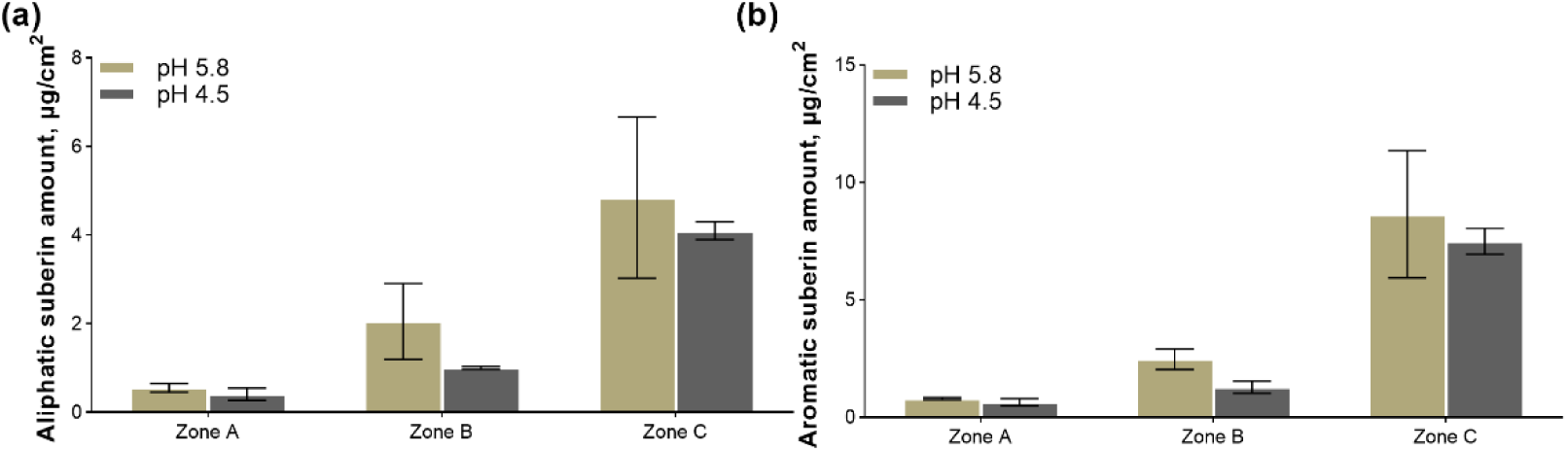
Total amounts of suberin in barley roots grown under different pH conditions. Total amounts of (a) aliphatic and (b) aromatic suberin in barley seminal roots grown under different pH conditions. Results are shown as mean expression ±SD of three biological replicates, no significant difference was detected.

**Supplemental Fig. S2.**
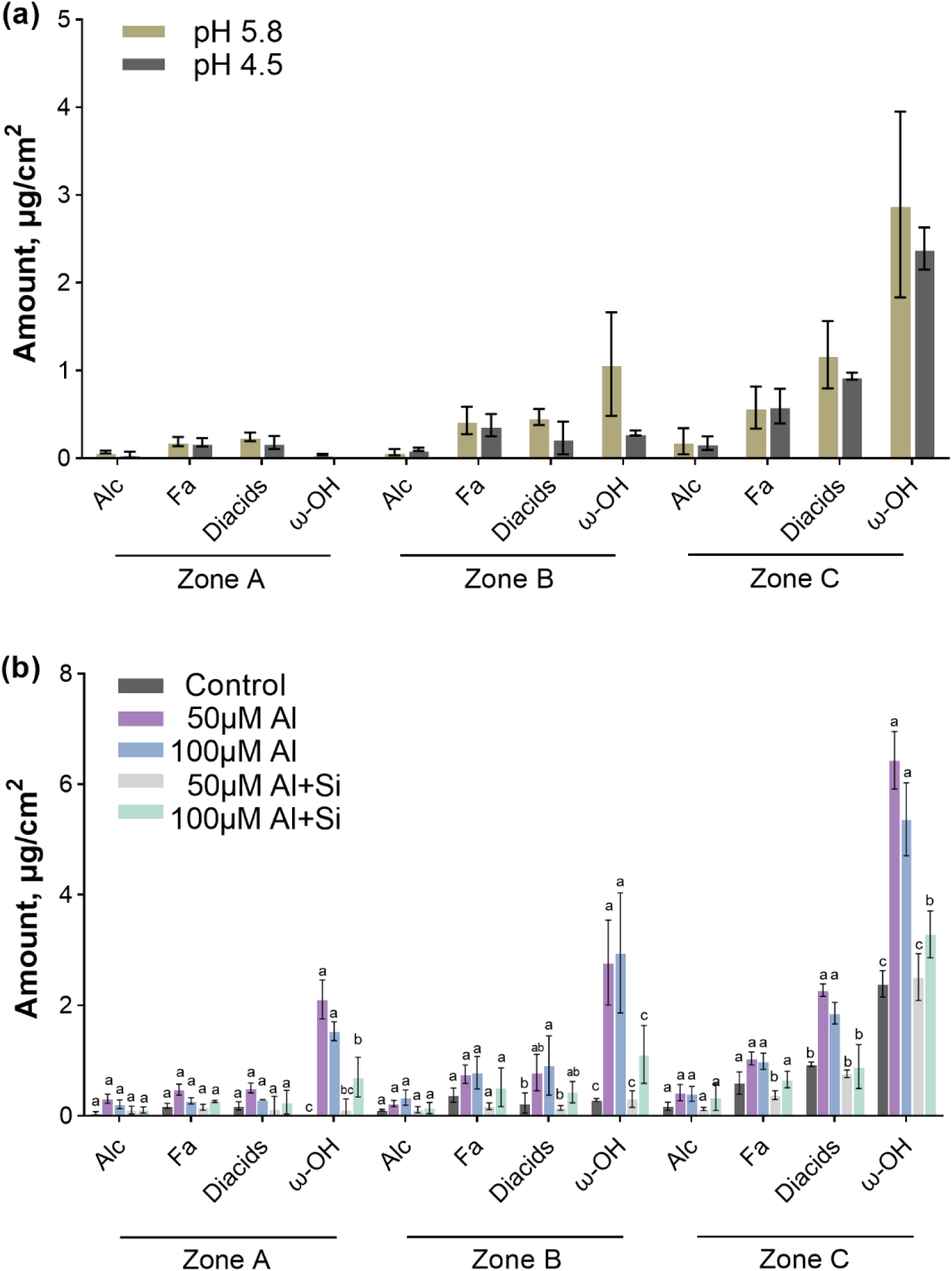
Amounts of substance classes of aliphatic suberin in barley seminal roots grown under different conditions. (a) Amounts of substance classes of aliphatic suberin in different zones of barley roots grown under different pH conditions and (b) under 50 μM or 100 μM Al treatment with or without Si conditions (at pH 4.5). Results are shown as mean expression ±SD of three biological replicates, different letters indicate significant differences (p< 0.05). No significant difference was detected in (a). Alc, primary alcohols; Fa, fatty acids; diacids, α–ω dicarboxylic acids; ω-OH acids, ω-hydroxy acids.

**Supplemental Fig. S3.**
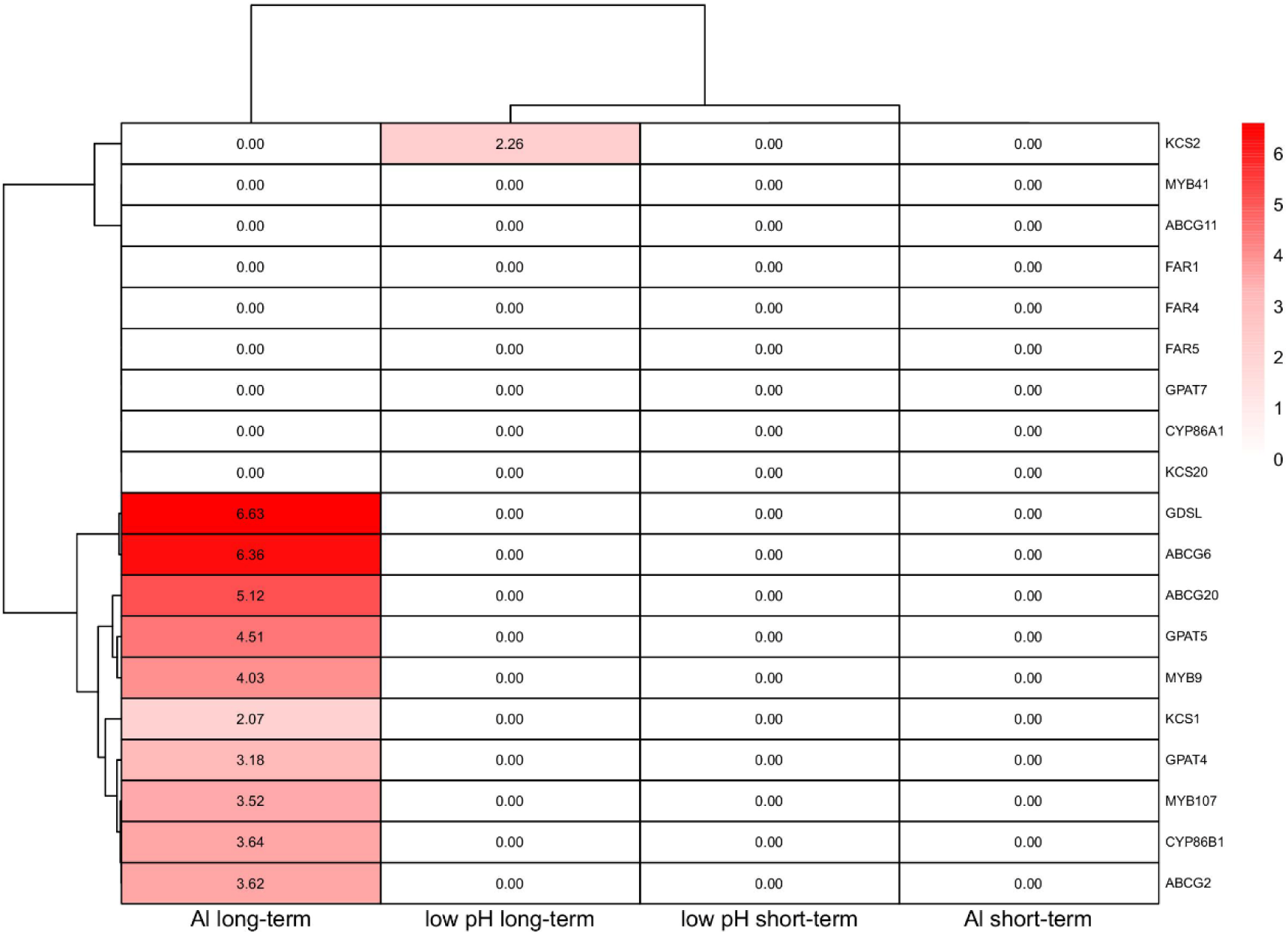
Expressions of selected suberin-biosynthesis genes of barley under different conditions. Expression data were obtained from (Szurman-Zubrzycka et al., 2021). The expression profile for each gene is shown as the log-ratio of signal intensity (log2 treat/control). Al long-term, 10 μM Al^3+^ for 7 days at pH 4.0; low pH long-term, 7 days at pH 4.0; low pH short-term, 24 h at pH 4.0; Al short-term, 10 μM Al^3+^ for 24 h at pH 4.0. control at pH 6.0.

**Supplemental Fig. S4.**
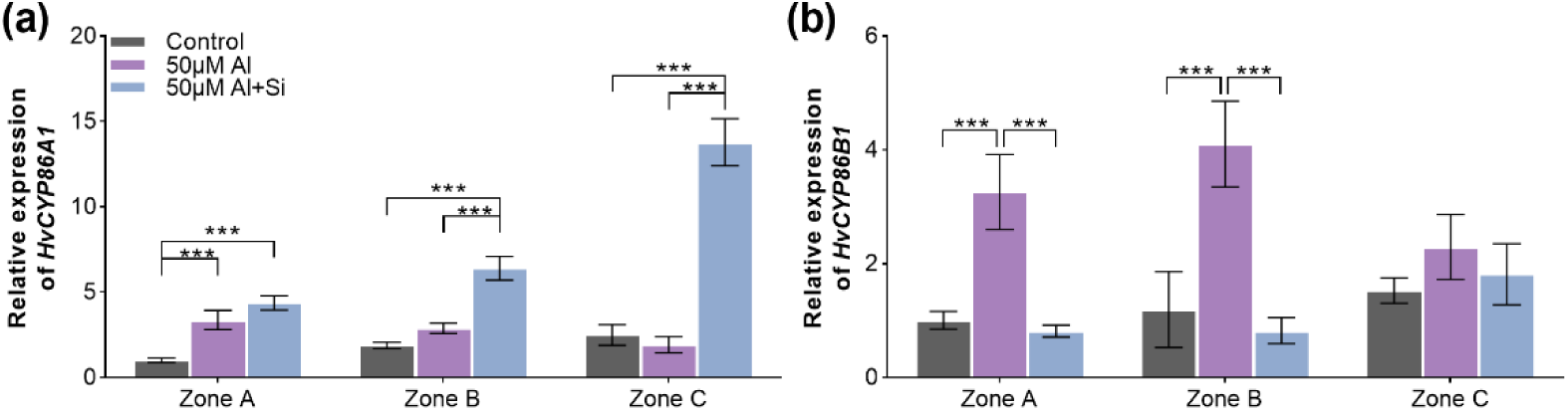
Expression analysis of *HvCYP86A1* and *HvCYP86B1* genes of barley using qPCR. Barley roots were exposed to a nutrient solution containing 0 M Al (control), 50 μM AlCl_3_ or 50 μM AlCl_3_ with 1mM Si additions for 4 days. Barley *ACTIN* and *GADPH* expression were used as a control. The gene expressions in zone A under control were arbitrarily fixed at 1. Results are shown as mean expression ±SD of three independent experiments, ***P < 0.001.

**Supplemental Fig. S5.**
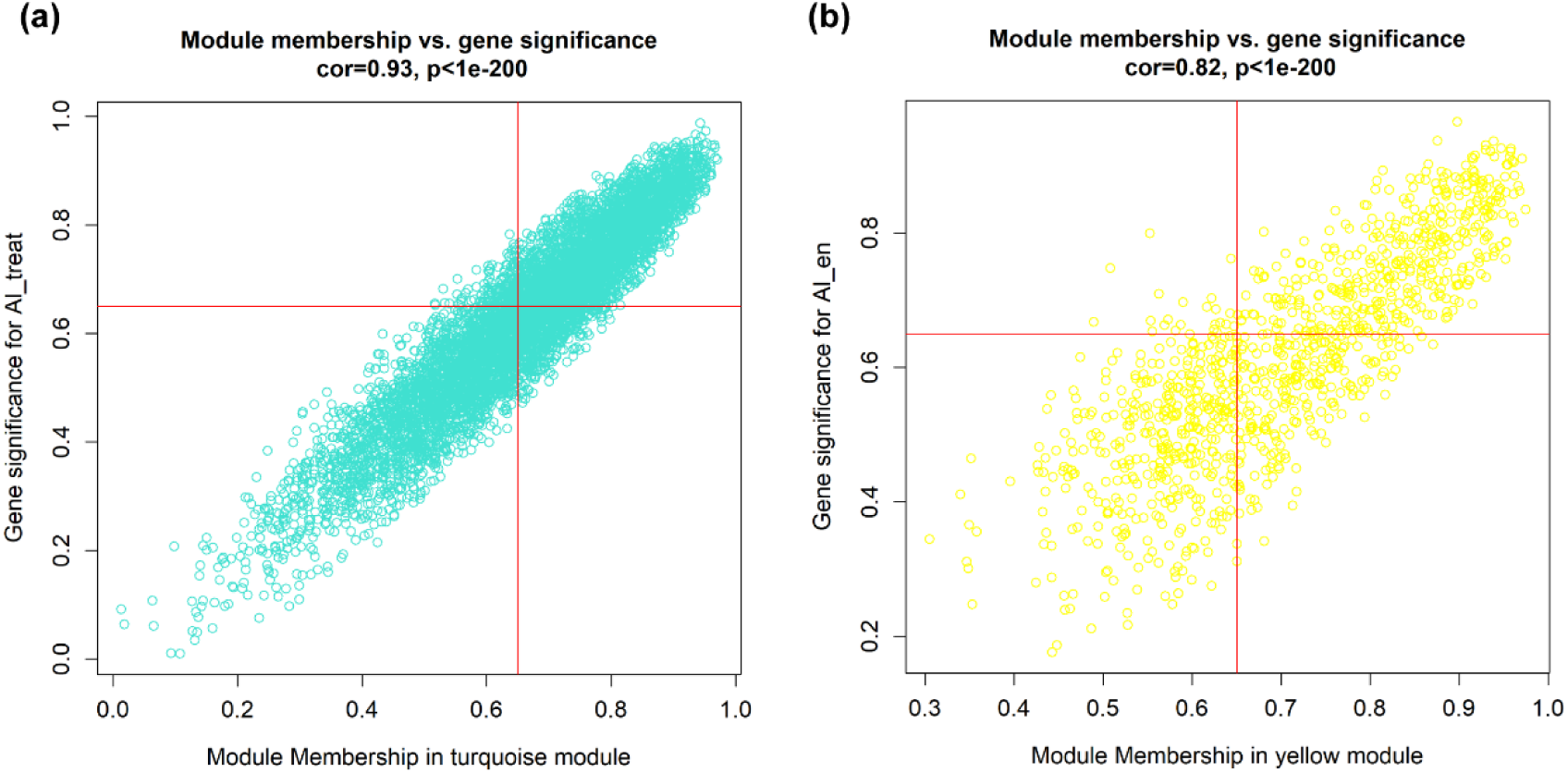
The correlation between Module Membership and Gene Significance. Analysis of Module Membership (MM) and Gene Significance (GS) in key co-expression modules under Al treatment (a) and endodermis under Al treatment (b). Scatter plots display the positive correlation between MM and GS for genes in the **MEturquoise** (a) and **MEyellow** modules (b). Each point represents a gene, with the red line indicating the threshold of 0.65 for both MM and GS. Genes above this threshold are considered highly association with the respective module and Al-related traits.

**Supplemental Fig. S6.**
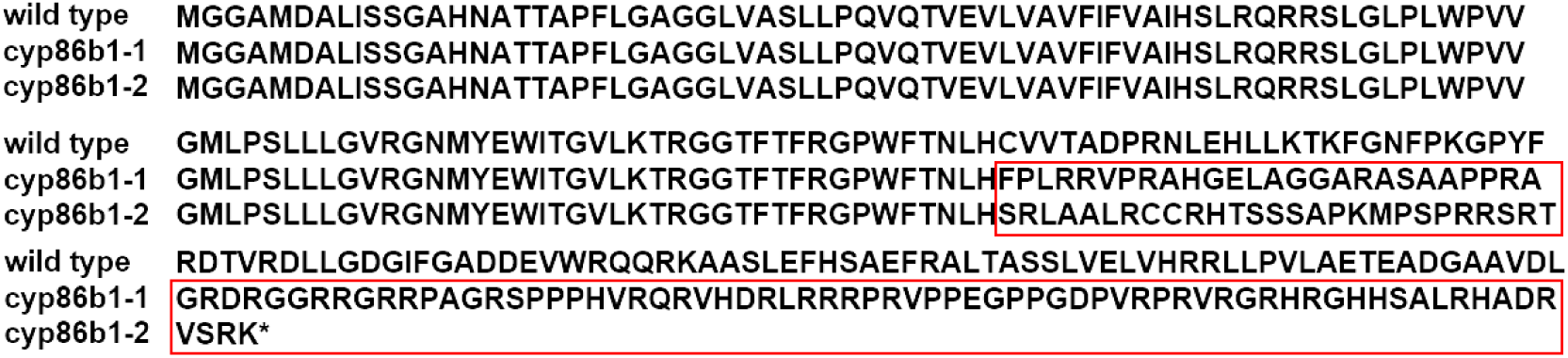
Protein sequence of *cyp86b1* mutants. Multiple sequence alignment of cyp86b1 protein. Red boxes mark the difference areas.

**Supplemental Fig. S7.**
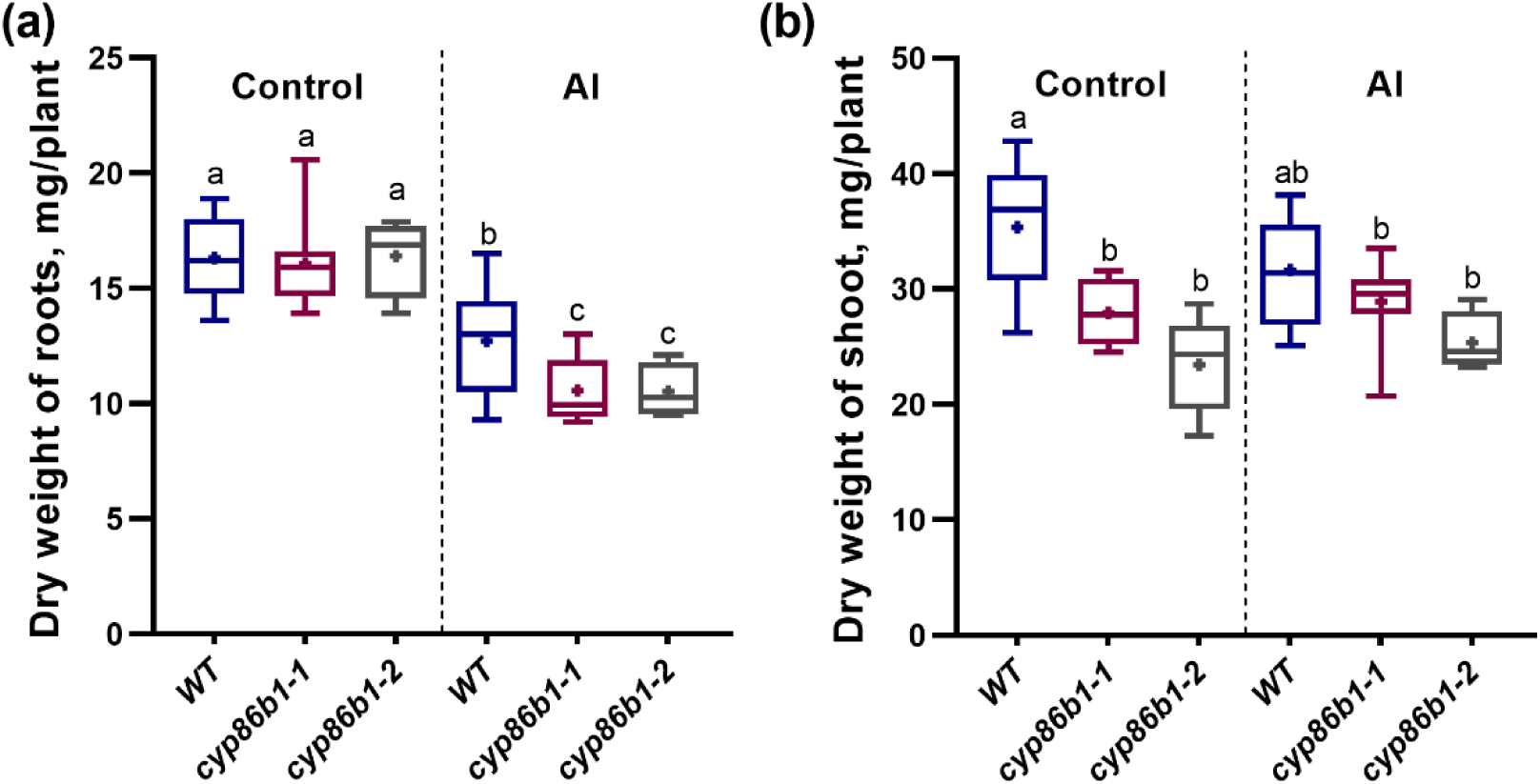
Effect of Al on shoot and root dry weight of wild type and transgenic barley plants. (a) Dry weight of roots and (b) shoots of 10-d-old GPF and barley mutants grown under different conditions. Barley plants grown in nutrient solution at pH 4.5 were presented as controls. The boxes range from the 10 to 90 percentiles. The “+” in the box represents the mean value. The whiskers range to the outliers. Different letters indicate significant differences (P < 0.05).

**Supplemental Fig. S8.**
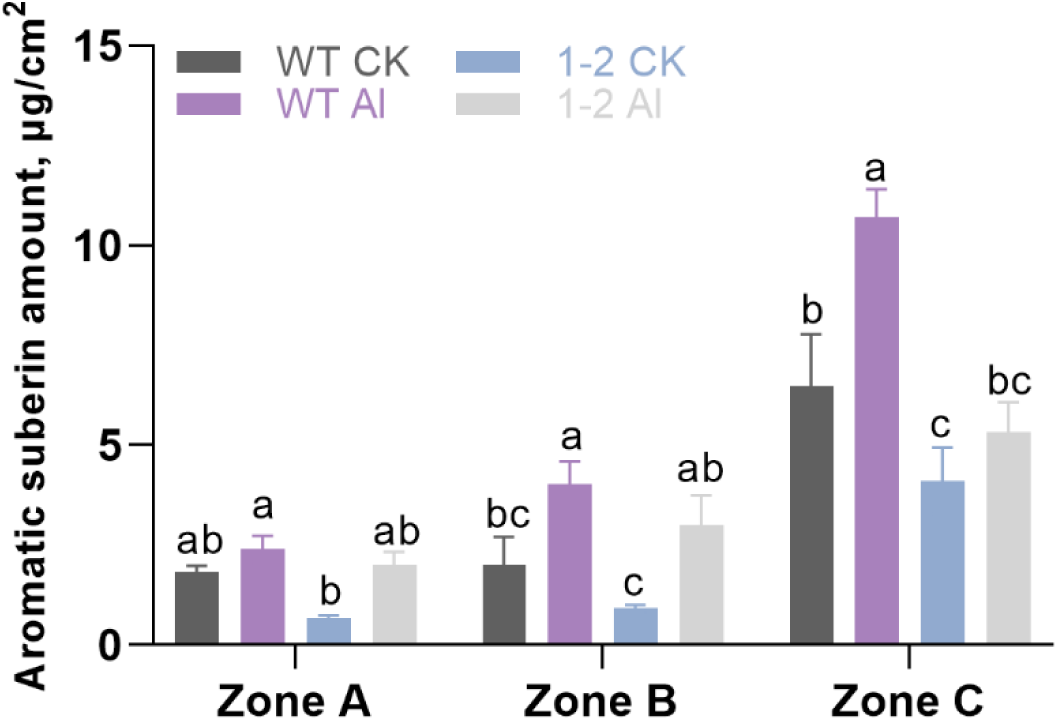
Aromatic suberin amount in barley mutant. Total amounts aromatic suberin in GPF and barley mutant seminal roots grown under different conditions. Results are shown as mean expression ±SD of three biological replicates, Different letters indicate significant differences (P < 0.05).

**Supplemental Fig. S9.**
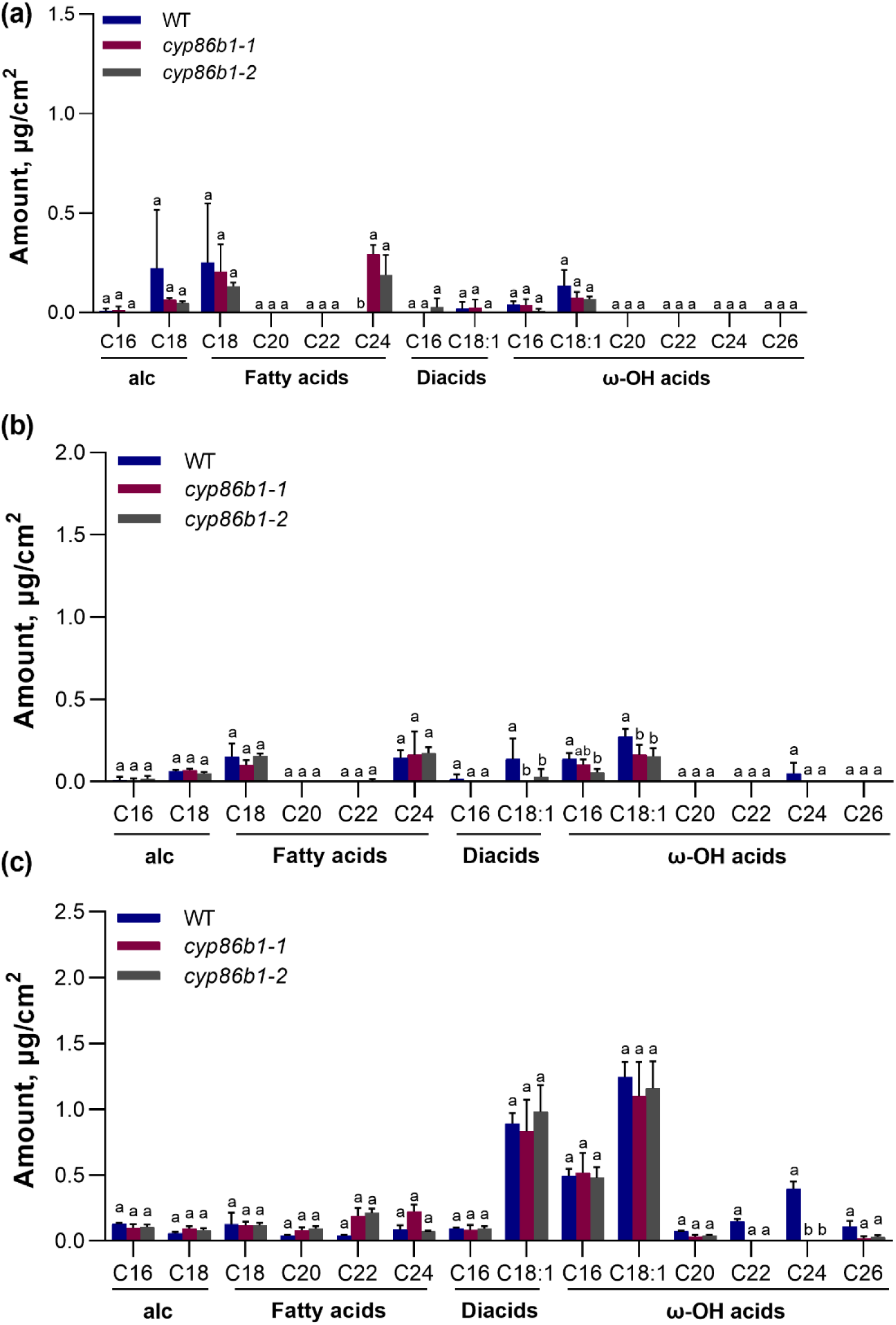
Amounts of monomers of aliphatic suberin in different zones of barley mutant roots. Aliphatic suberin monomers amount of barley (GPF) roots zone A (a), zone B (b), zone C (c). Plants grown in nutrient solutions at pH 4.5. Results are shown as mean expression ±SD of three biological replicates, different letters indicate significant differences (p< 0.05). alc, primary alcohols; diacids, α–ω dicarboxylic acids; ω-OH acids, ω-hydroxy acids.

**Supplemental Fig. S10.**
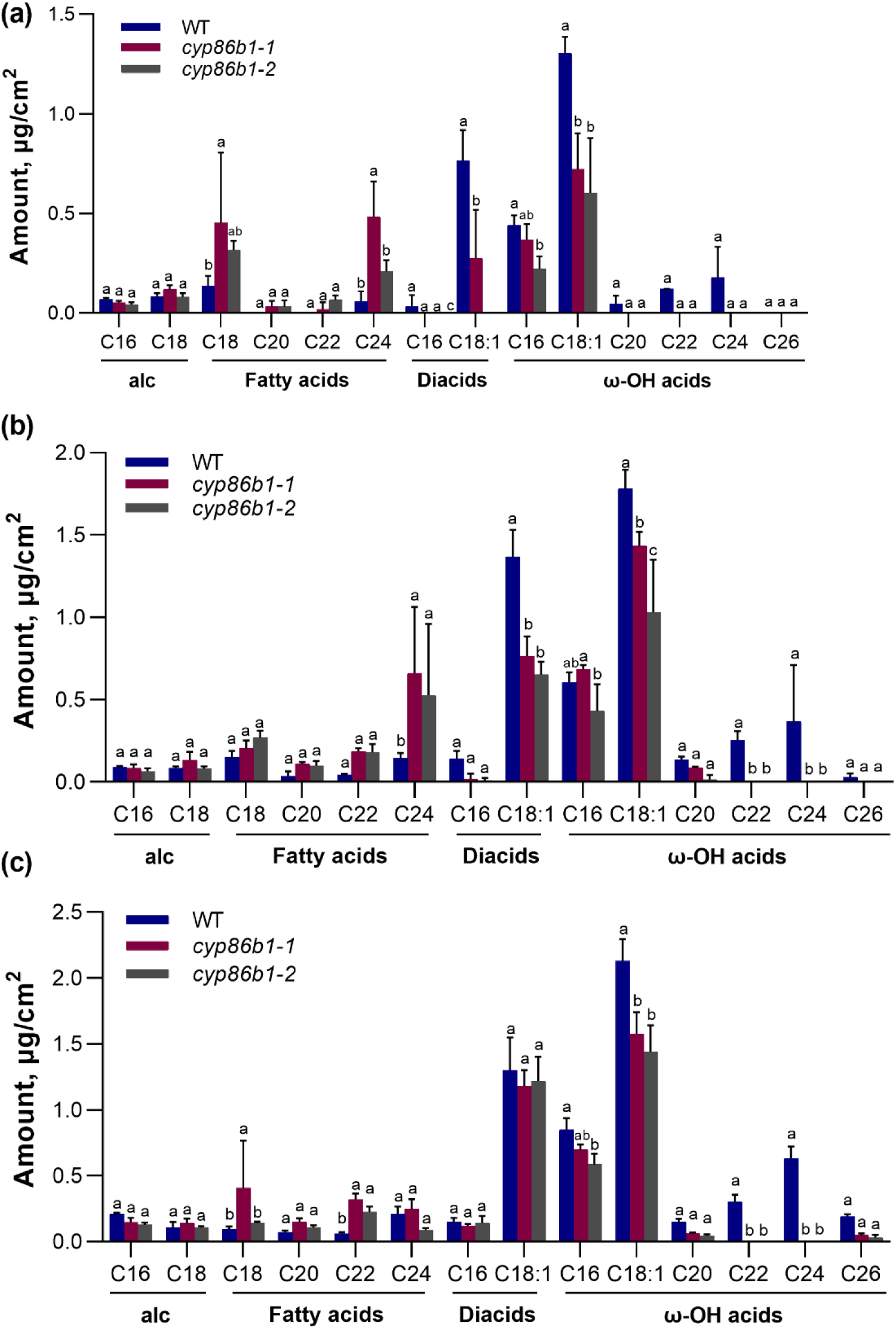
Amounts of monomers of aliphatic suberin in different zones of barley mutant roots under Al conditions. Aliphatic suberin monomers amount of barley (GPF) roots zone A (a), zone B (b), zone C (c). Plants grown in nutrient solutions at pH 4.5 with 50 μM AlCl_3_. Results are shown as mean expression ±SD of three biological replicates, different letters indicate significant differences (p< 0.05). alc, primary alcohols; diacids, α–ω dicarboxylic acids; ω-OH acids, ω-hydroxy acids.

**Supplemental Fig. S11.**
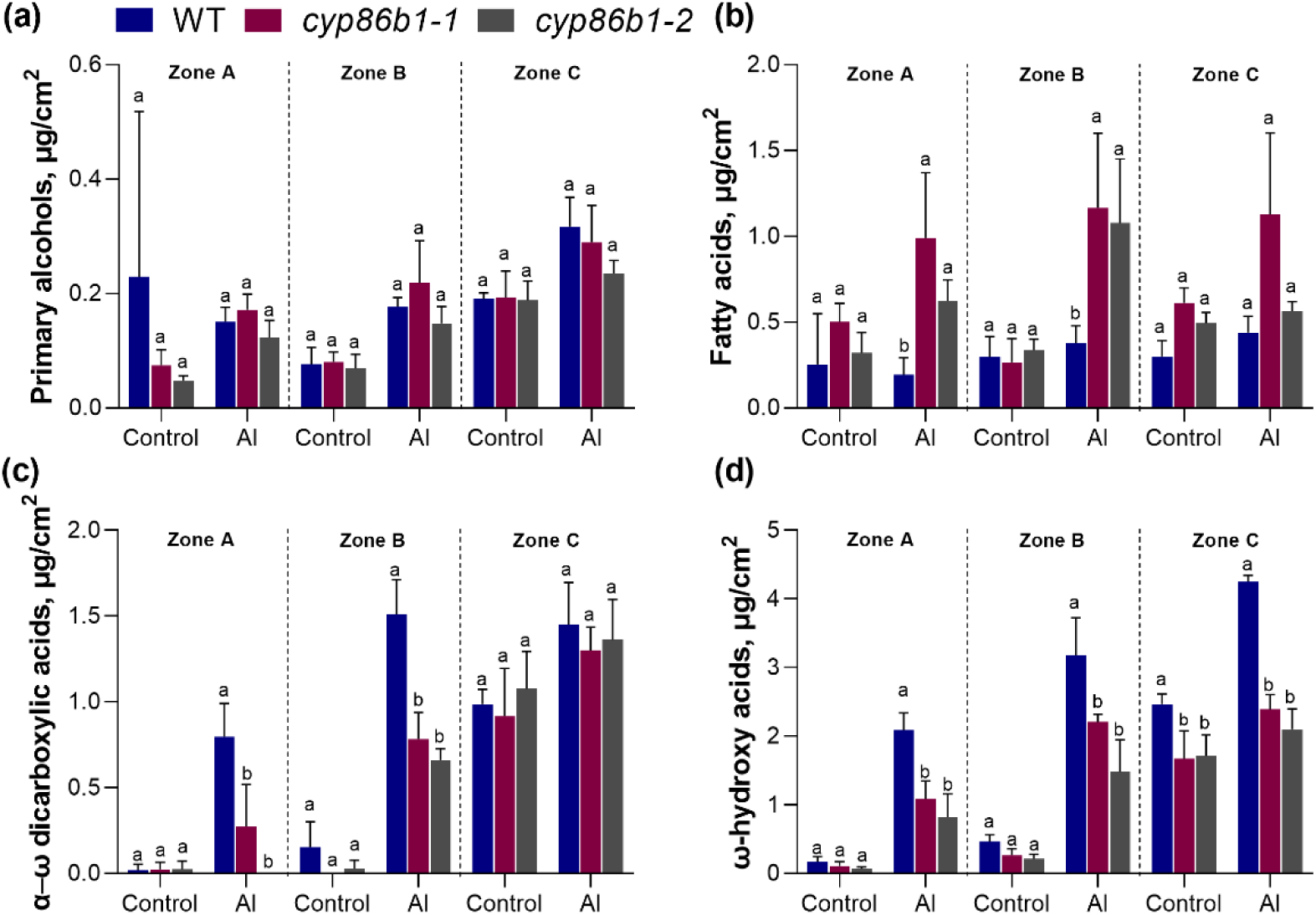
Amounts of substance classes of aliphatic suberin in wild type and transgenic barley seminal roots. Amounts of (a) primary alcohols, (b) fatty acids, (c) α–ω dicarboxylic acids, and (d) ω-hydroxy acids in different zones of barley roots grown under different. Results are shown as mean expression ±SD of three biological replicates, different letters indicate significant differences (p< 0.05).

**Supplemental Fig. S12.**
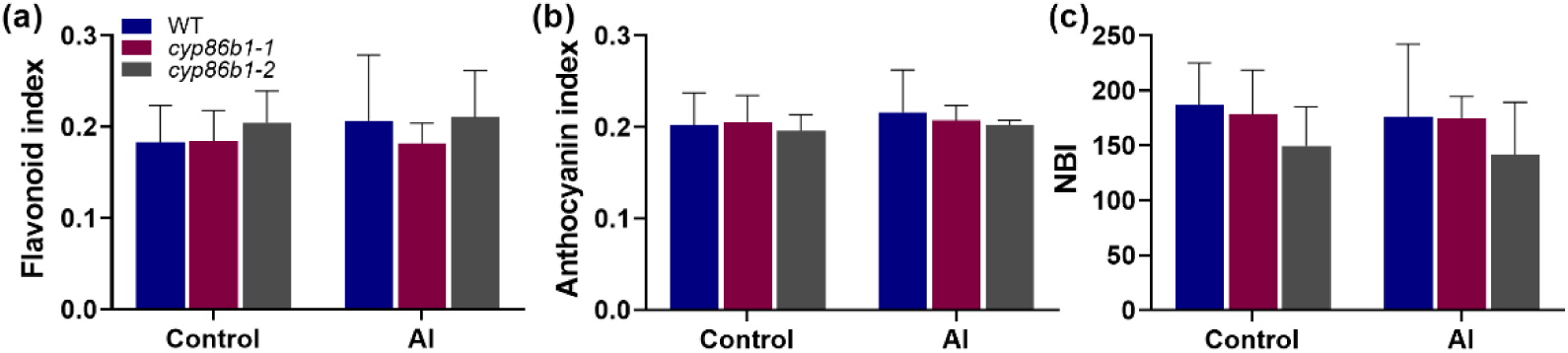
Effect of Al stress on Flavonoid index, Anthocyanin index, Nitrogen Balance Index of GPF and barley mutants. (a) Flavonoid index, (b) Anthocyanin index, and (c) Nitrogen Balance Index (NBI) of GPF and barley mutants grown under different. Results are shown as mean expression ±SD. No significant difference was detected.

**Supplemental Fig. S13.**
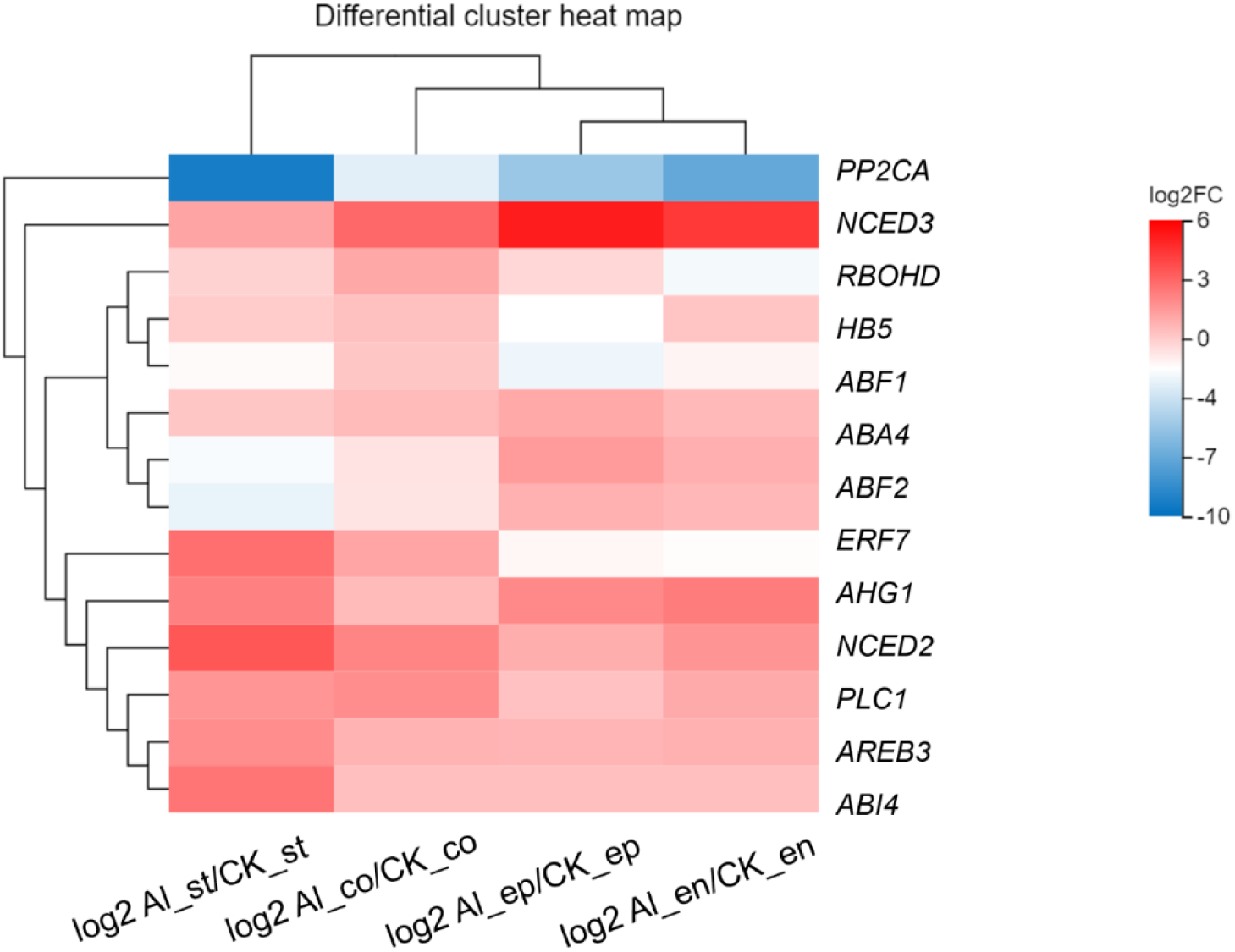
Expressions of selected ABA pathway genes of barley under different conditions. Heatmap of log_2_ fold change (log2 Al /CK) of selected ABA pathway genes in four different tissues. Red represents up-regulated, blue represents down-regulated. ep, epidermis; co, cortex; en, endodermis; st, stele.

**Supplemental Fig. S14.**
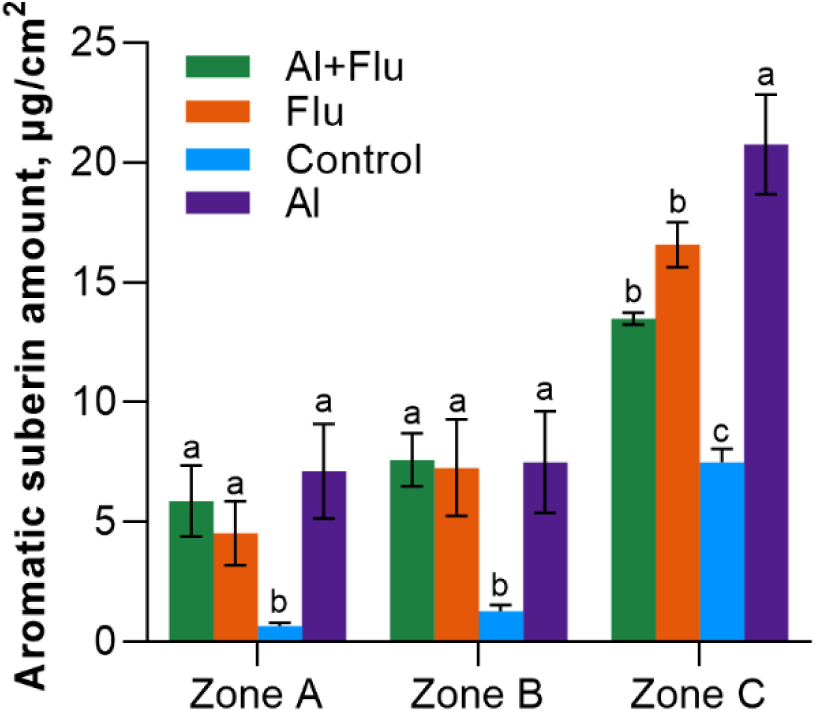
Aromatic suberin amount in barley roots under Flu treatment. Total amounts aromatic suberin in barley (cv. Scarlett) seminal roots grown under different conditions. Results are shown as mean expression ±SD of three biological replicates, Different letters indicate significant differences (P < 0.05).

